# A tissue-resolved transcriptomic atlas of adult male *Halyomorpha halys* reveals tissue-specific RNAi machinery and a minimal systemic response to non-specific dsRNA

**DOI:** 10.64898/2026.05.26.728018

**Authors:** Venkata Partha Sarathi Amineni, Swathi Ramapuram, Kristen A. Panfilio

## Abstract

**Background:** *Halyomorpha halys* (brown marmorated stink bug) is an invasive polyphagous pest causing significant agricultural damage worldwide and is an emerging target for RNAi-based pest management. Despite growing interest in dsRNA-based biocontrol, progress is constrained by the lack of tissue-resolved transcriptomic resources covering key biological processes such as feeding, detoxification, and reproduction. Furthermore, our understanding of how RNAi machinery expression varies across tissues remains limited, which impairs both target gene selection and predictions of RNAi efficacy. Critically, the transcriptional response of *H. halys* to haemolymph-delivered non-specific dsRNA represents a key knowledge gap for evaluating potential non-target immune reactions of dsRNA-based approaches.

**Results:** Field-collected adult males were injected with either nuclease-free water or dsRNA targeting GFP (dsGFP), and transcriptomes were generated from the brain, midgut, salivary glands, and testes. Sequencing produced high-quality datasets with clear tissue-level separation and tight clustering of biological replicates. As expected in targeting a non-endogenous gene, differential expression analysis revealed a limited transcriptional response to dsGFP. Baseline profiling of RNAi pathway genes in controls showed broad expression of core siRNA and miRNA components across all tissues, yet with marked specialisation: two additional Argonaute-2 isoforms and multiple piRNA factors were testes-specific, whereas salivary glands showed strong, restricted expression of nuclease-encoding genes, including a T2 ribonuclease and a non-specific endonuclease. Expression atlases also revealed pronounced tissue partitioning for other protein families. Consistent with their respective functions, secreted trypsins and chymotrypsins are salivary-enriched while the cathepsins for intracellular protein catabolism are midgut-enriched, with brain-centred neuropeptide expression. However, we also uncovered unexpected nuance, such as closely related subfamilies of Cytochrome P450s, which generally function as detoxification enzymes, being partitioned between the midgut, brain or testes.

**Conclusions:** This work delivers the first tissue-resolved transcriptomic atlas of adult male *H. halys*, providing a high-resolution resource on compartmentalization of proteolysis, detoxification, and neuroendocrine signalling, as well as for candidate gene discovery in RNAi-based pest control. The modest, tissue-restricted transcriptional response to non-specific dsRNA, together with strong tissue-specific enrichment of some components, offers mechanistic insight into tissue-dependent RNAi efficiency and supports rational dsRNA target selection in *H. halys*.

## Introduction

The brown marmorated stink bug, *Halyomorpha halys*, is a highly polyphagous hemipteran pest that causes substantial damage to fruit, vegetable, and field crops worldwide [1–3], with a growing need for genomics approaches to characterise invasive populations [4]. Its success as a pest is linked to high fecundity, a broad host range, and a piercing–sucking feeding mode in which secreted saliva injures plant tissues, leading to necrotic lesions and malformed fruit that reduce the marketable value of the produce [3]. Together, these features make *H. halys* a persistent challenge for sustainable pest management.

In recent years, genomic and transcriptomic resources for *H. halys* have expanded considerably. A 1.15-Gb draft genome generated under the i5k initiative provides a well-annotated reference that highlights gene families associated with sensory function, digestion, immunity, detoxification, and development [5–7]. Chemosensation, focusing on olfaction and odorant-binding proteins, has been well characterised in antennal transcriptomes [8, 9]. Whole-body and developmental transcriptomes spanning multiple life stages and both sexes further describe global gene expression dynamics and have been used to characterize candidate genes across functional categories, including detoxification and stress responses [6, 10]. Proteomic and tissue-specific transcriptomic studies of salivary glands and the gut have revealed that watery and sheath saliva comprise distinct protein suites, and that the principal salivary gland is a major source of digestive proteases and nucleases [11, 12]. While these studies provide valuable insight into specific organs and biological processes, a multi-tissue, gene-level framework remains lacking.

RNA interference (RNAi) has emerged as a promising strategy for species-specific pest control, and *H. halys* is known to be RNAi-competent when dsRNA is delivered by injection [13–15]. However, oral delivery of dsRNA has proven much less effective [14]. In our previous work, we showed that *H. halys* saliva exhibits strong dsRNase activity that rapidly degrades dsRNA and that co-formulation with double-stranded DNA can partially protect dsRNA and enhance gene silencing, although this did not fully translate into increased mortality [14]. These findings indicate that nuclease activity in the oral cavity is a major barrier to RNAi but also suggest that additional factors, such as tissue-specific uptake, intracellular processing, and systemic spread, could likely influence RNAi efficiency in this species [16, 17].

Despite the growing interest in RNAi-based control, our understanding of how RNAi-related gene activity is organised across tissues in *H. halys* remains limited. In particular, the tissue-specific expression of key RNAi pathway components, including Dicers, Argonautes, transporters, and nucleases, has not been systematically characterised across major organs. Likewise, although detoxification enzymes such as cytochrome P450s, glutathione S-transferases, and carboxylesterases are known to be expanded in *H. halys* and contribute to its polyphagy and xenobiotic tolerance [18], their distribution across tissues in a unified transcriptomic framework remains incompletely resolved. Neuropeptides and their G protein-coupled receptors (GPCRs), which are increasingly recognised as attractive targets for precision pest control [19, 20], have also not been comprehensively mapped across different individual tissues in this species [21, 22].

In addition to baseline expression patterns, it remains unclear how gene expression responds across tissues following exposure to dsRNA in *H. halys*. While RNAi is often associated with antiviral and immune-like responses in some insect systems, it is not known whether haemolymph-delivered dsRNA induces coordinated transcriptional changes across tissues in *H. halys*, or whether such responses are limited and tissue-specific [23–27]. Similar to baseline transcriptomic studies in other insects, it is also important to establish the extent to which dsRNA targeting a non-endogenous sequence like *gfp* affects global gene expression in a given species [28–30]. Addressing this question is important for understanding both the RNAi mechanism and the potential for unintended transcriptional effects following dsRNA exposure.

To address these gaps, we generated a tissue-resolved RNA-seq atlas of adult male *H. halys*, focusing on four functionally distinct tissues: brain (central nervous system), midgut (excluding the posterior midgut with enriched symbiont), salivary glands, and testes. Using a control injection of non-specific dsRNA (dsGFP) alongside water-injected controls, we further assessed the transcriptional response to dsRNA exposure at 72 h post-injection. Gene-level expression was quantified using a genome-guided alignment and normalisation framework to enable robust cross-tissue comparisons.

This study was designed to (i) characterise baseline tissue-specific expression of RNAi pathway components, nucleases, and key functional gene families, (ii) identify tissue-specific enrichment patterns relevant to RNAi efficacy, and (iii) evaluate the extent and tissue distribution of transcriptional responses to dsRNA exposure. In addition, by integrating tissue-specificity metrics with expression magnitude, we identify highly expressed, tissue-restricted genes that provide a rational basis for prioritising candidate RNAi targets, as these genes are likely to reflect core functional processes within individual tissues and may enable more targeted and effective gene silencing strategies.

By integrating these analyses, we provide a comprehensive tissue-level resource for *H. halys* and a framework for understanding tissue-dependent RNAi efficiency, while also enabling broader insights into tissue-specific gene regulation. Together, these findings have direct implications for the rational development of dsRNA-based pest management strategies in *H. halys*.

## Methods

### Insects, tissue collection, and experimental design

Adult male *Halyomorpha halys* were collected from *Catalpa spp*. trees in the botanical garden of the University of Hohenheim, Stuttgart, Germany (48.707561° N, 9.211532° E) during September–October 2025. Insects were maintained under controlled laboratory conditions (25°C, 65% relative humidity, 16:8 h light:dark photoperiod) for 5–15 days prior to treatment, on a diet of *Catalpa spp.* seed pods from the host plant and pesticide-free apples supplemented with tap water provided via a cotton-plugged falcon tube.

For the injection experiment, insects were anaesthetised on ice and received a ventral abdominal injection of either 2 µg of double-stranded RNA targeting the *green fluorescent protein* coding sequence (dsRNA-GFP sequence available in supplementary file SI_1; synthesized by Genolution Inc., Seoul, Republic of Korea) dissolved in 2 µL nuclease-free water, or 2 µL nuclease-free water alone (untreated control, hereafter UNT). Tissues were collected 72 h post-injection. Prior to dissection, insects were anaesthetised on ice for 5–10 min and dissected in ice-cold 1× phosphate-buffered saline (PBS). Four tissues were collected per individual: brain/CNS, midgut (the posterior, red-coloured symbiont-enriched region was excluded), salivary glands, and testes (Fig. 1). Tissues from five adult males were pooled per biological replicate and snap-frozen immediately in liquid nitrogen. Three biological replicates were generated per tissue per treatment condition, yielding 24 libraries in total (4 tissues × 2 treatments × 3 replicates). For the tissue-resolved expression atlas and all gene-family profiling analyses reported here, only the water-injected control libraries were used (n = 3 biological replicates per tissue; 12 libraries total).

**Figure 1.**
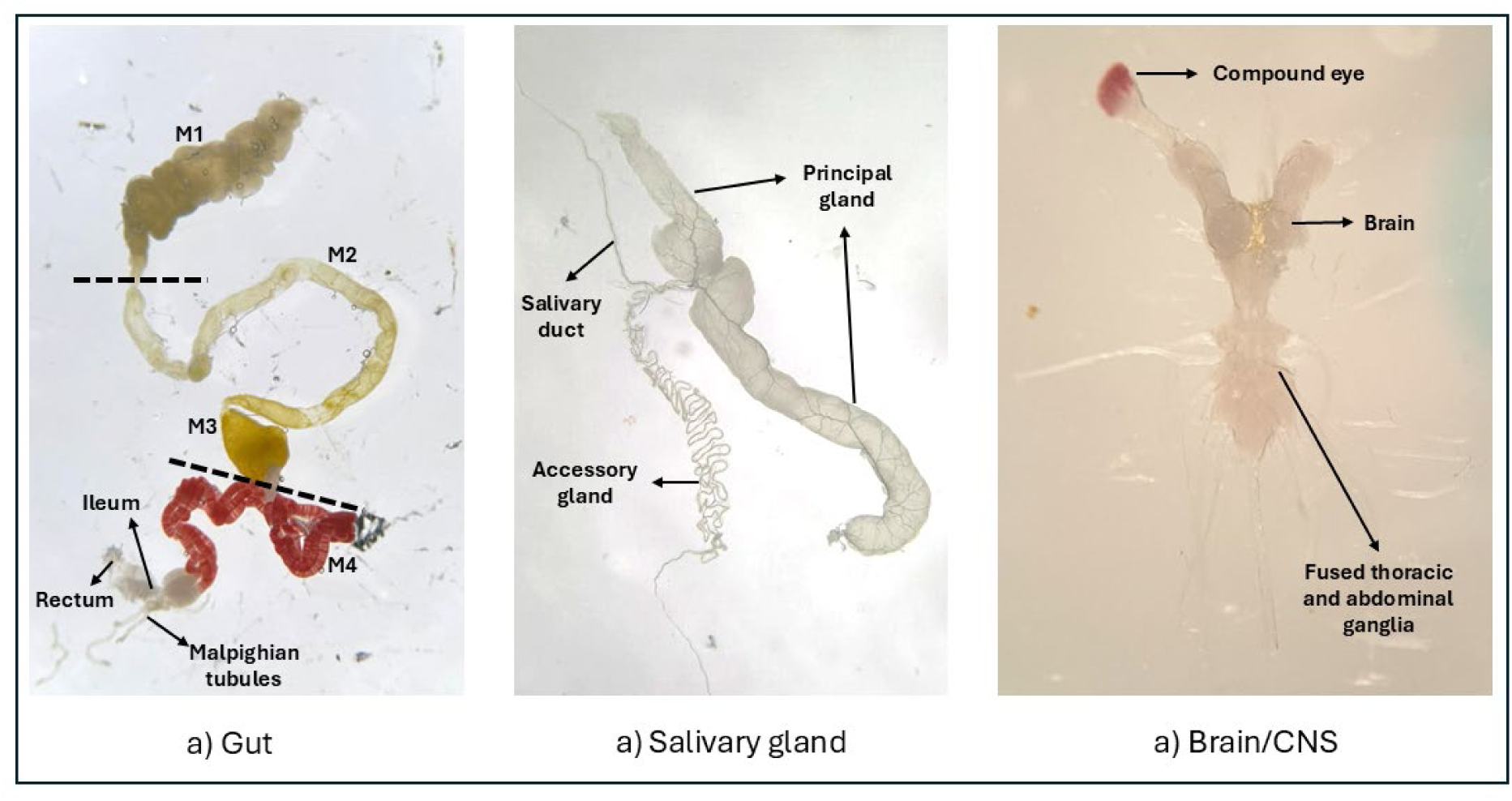
Dissection of tissues used for transcriptomic analysis in Halyomorpha halys. Representative images of dissected tissues: (a) gut, (b) salivary glands, and (c) brain/CNS. In (a), the midgut is shown with regions labelled M1–M4, along with the ileum, rectum, and Malpighian tubules. The dotted lines indicate the midgut region (M2 & M3) that was dissected and used for downstream analyses. M = midgut. In (b), the salivary gland complex includes the principal gland, accessory gland, and salivary duct. In (c), the central nervous system shows the brain (supraesophageal ganglion), compound eye (peripheral nervous system), and the fused thoracic and abdominal ganglia, connected via the ventral nerve cord; all tissues shown were used for RNA isolation in the experiments.

### RNA extraction, library preparation, and sequencing

Total RNA was extracted from each pooled tissue sample using the NEB Total RNA Preparation Kit (New England Biolabs, Heidelberg, Germany) with on-column DNase I treatment according to the manufacturer’s protocol. Extracted RNA was submitted to Novogene GmbH (Munich, Germany) for polyA-enriched library preparation and 150-bp paired-end Illumina sequencing (PE150). Only samples with an RNA Integrity Number (RIN) ≥ 7 were chosen and proceeded to library construction. Sequencing output ranged from approximately 9.2 to 16.5 Gb per sample (∼30.5–55 million read pairs per library).

### Read quality control, genome alignment, and gene-level quantification

Raw sequencing data were retrieved from compressed archives and file integrity verified using MD5 checksums. Where technical lane replicates were present, paired-end FASTQ files were merged per sample prior to downstream processing. Initial read quality was assessed using FastQC and MultiQC [31, 32]. Adapter sequences and low-quality bases were removed using fastp (v1.0.1) with paired-end adapter auto-detection, poly-G and poly-X tail trimming, a minimum mean base quality of Q30, and a minimum post-trimming read length of 50 bp [33]. Post-trimming quality was re-assessed with FastQC and MultiQC to confirm adequate filtering.

Genome alignment and gene-level quantification were performed on the bwHPC high-performance computing cluster (Baden-Württemberg High Performance Computing, Germany). Trimmed reads were aligned to the *Halyomorpha halys* reference genome assembly (GCF_000696795.3-RS_2024_07, accessed via NCBI RefSeq) using STAR (v2.7.11b) in two-pass mode (--twopassMode Basic). Alignments were output as coordinate-sorted BAM files, permitting a maximum of 20 multiple alignments per read (--outFilterMultimapNmax 20) and up to 10 mismatches per read (--outFilterMismatchNmax 10). Alignment summary statistics, including uniquely mapped and multi-mapping read fractions, were extracted from STAR log files (Log.final.out) and aggregated across all 24 libraries.

Gene-level read counts were generated from coordinate-sorted BAM files using HTSeq-count (v2.0.9) in union mode (-m union), with position-sorted alignments (-r pos), unstranded library settings (-s no), exon features (-t exon), and gene identifiers (-i gene_id) from the reference GTF annotation (NCBI assembly: GCF_000696795.3-RS_2024_07). The GTF annotation contains 17,693 genes, of which exon coordinates are available for 17,483 genes. Secondary and supplementary alignments were excluded from quantification (--secondary-alignments ignore, --supplementary-alignments ignore). Only uniquely mapping primary alignments were retained; reads overlapping multiple annotated genes or mapping to multiple genomic loci were discarded, yielding a conservative count matrix of unambiguously assignable reads for all downstream analyses.

During gene-family expression profiling, one brain biological replicate from each treatment group (brain replicate 1, UNT and brain replicate 1, dsGFP) was identified as exhibiting anomalous expression patterns within the nuclease and protease gene sets, with expression profiles systematically inconsistent with the remaining brain replicates. As this deviation was not detectable by PCA of global variance-stabilised counts, it was identified through targeted inspection of normalised expression values within specific gene families. These two samples were excluded from all downstream analyses. See Fig. S1 in supplementary data file SI_1 for further details. Accordingly, all brain-tissue analyses were conducted using n = 2 biological replicates per treatment group, while midgut, salivary glands, and testes retained n = 3 biological replicates per treatment group.

### Differential expression analysis

Differential gene expression between dsRNA-GFP-injected and water-injected samples was assessed using DESeq2 (v1.42.1) implemented in R (v4.3.3; 2024-02-29) running under Windows Subsystem for Linux (WSL) [34]. Raw HTSeq-count matrices were used as input [35], with size-factor normalisation performed within DESeq2 prior to fitting a negative binomial model. Analyses were conducted independently for each tissue using the design formula ∼ treatment. Prior to statistical testing, genes with counts of at least 5 in at least 3 samples, or in all samples when fewer than 3 samples were available in an analysis set, were retained for analysis. Differential expression significance was assessed using the Benjamini–Hochberg procedure to control the false discovery rate (FDR), with a significance threshold of adjusted p < 0.05.

### Cross-tissue expression profiling, normalisation, and expression filtering

To construct a tissue expression atlas independent of treatment effects, only water-injected control samples were used (n = 3 biological replicates for testes, salivary gland, midgut, and n=2 for brain; 11 libraries total). A multi-tissue DESeq2 object was constructed with the design formula ∼ tissue, and size-factor normalisation was applied to the full count matrix. Normalised expression values were log₂-transformed as log₂(normalised count + 1) for cross-tissue comparisons and visualisation. For tissue-level summaries, mean log₂-transformed values were calculated across the three biological replicates within each tissue.

To reduce noise in gene-family visualisations, a minimum expression threshold was applied: genes were retained for display only if their mean normalised count was ≥ 10 in at least one tissue. This filtering was applied exclusively during heatmap visualisation and did not influence any differential expression analysis. Tissue enrichment patterns were characterised using both absolute expression values and row-wise z-score standardisation, where each gene’s expression was centred and scaled to unit variance across the four tissues.

### Heatmap visualisation

Heatmaps were generated in R (v4.3.3) using the *pheatmap* package (v1.0.12). Two complementary approaches were used: (i) absolute expression heatmaps based on log₂(normalised count + 1) values, representing expression magnitude; and (ii) row-scaled heatmaps in which expression values were z-score standardised per gene, highlighting relative tissue enrichment independently of absolute expression level. Tissue-level heatmaps used mean normalised expression values per tissue. Rows were hierarchically clustered using Euclidean distance with complete linkage; columns were ordered by tissue identity and replicate number without clustering.

### Gene-family expression profiling

Candidate gene sets for neuropeptides, neuropeptide receptors, detoxification enzymes (cytochrome P450s, glutathione S-transferases, carboxyl/hormone esterases), nucleases, proteases, aminopeptidases, carboxypeptidases, and RNAi pathway components were compiled from published pentatomid transcriptomic resources and functional annotation studies [10–12, 18, 21, 36]. Gene identifiers were harmonised to the *H. halys* RefSeq annotation (GCF_000696795.3) using curated annotation tables cross-referenced against published gene lists. For each gene family, normalised expression matrices were extracted from the control-only dataset and visualised using log₂(normalised count + 1) values in separate heatmaps scaled dynamically by gene set size. Both absolute and z-score heatmaps were used to distinguish expression magnitude from tissue-specific enrichment patterns.

### Neuropeptide–receptor network analysis

Genes annotated as neuropeptides and receptors were assigned to functional classes using keyword-based classification of gene annotations. Tissue-enriched genes were identified across brain, midgut, salivary glands, and testes using a row-wise z-score threshold of ≥1.0 combined with an absolute expression threshold of log2 normalised count ≥3.0. For each tissue, co-enriched neuropeptide and receptor classes were summarised into class-level association networks, where edge weights were proportional to the number of genes supporting each class pairing.

### Tissue-specificity analysis (Tau)

Gene expression specificity across the four profiled tissues was quantified using the Tau (τ) metric [37, 38]. Raw HTSeq-count matrices were first normalised to counts per million (CPM) to correct for library size differences. For each gene, mean CPM values were calculated across the three biological replicates within each tissue (except n=2 in brain tissue), using control samples only. Tau was calculated per gene from the four-tissue mean expression vector, where τ = 0 indicates uniformly broad expression and τ = 1 indicates expression confined to a single tissue. Genes were classified as tissue-specific (τ ≥ 0.85), broadly expressed (τ ≤ 0.15), or intermediate. For expression breadth assignment, a gene was considered expressed in a given tissue only if it met both an absolute threshold (CPM ≥10) and a relative threshold (expression ≥5% of the gene’s maximum CPM across tissues). Genes were subsequently categorised as single-tissue expressed, multi-tissue expressed or ubiquitously expressed. The tissue with the highest mean CPM was designated the dominant tissue; genes satisfying the expression threshold in only one tissue were additionally classified as strictly tissue-specific.

## Results

### A high quality, tissue-specific transcriptomic resource for adult male *H. halys*

A total of 24 RNA-seq libraries were generated from four tissues (brain, midgut, salivary glands, and testes), with three biological replicates each for dsGFP-injected and water-injected adult male *H. halys*. Sequencing yielded high-quality paired-end reads (PE150) with consistently low error rates (0.01%) and high base-call quality scores (Q30 ≥95.9–98.1% per library). Raw library sizes ranged from approximately 30 to 55 million reads per sample (paired-end), providing sufficient sequencing depth for robust differential expression and cross-tissue profiling analyses. Adapter trimming and quality filtering retained more than 96% of reads across all tissues, with post-trimming Q30 scores of 96.87–98.44%.

In mapping to the genome assembly, uniquely mapped reads ranged from 85.7% to 97.1% across all libraries, and multi-mapping rates ranged from 0.99% to 9.02% (highest multi-mapping rate for the salivary gland, lowest for the testes). These results indicate consistently high mapping efficiency and low ambiguity in read assignment across tissues (Fig. 2A; SI_2). The complete count data table is provided in the supplementary information (SI_10).

**Figure 2.**
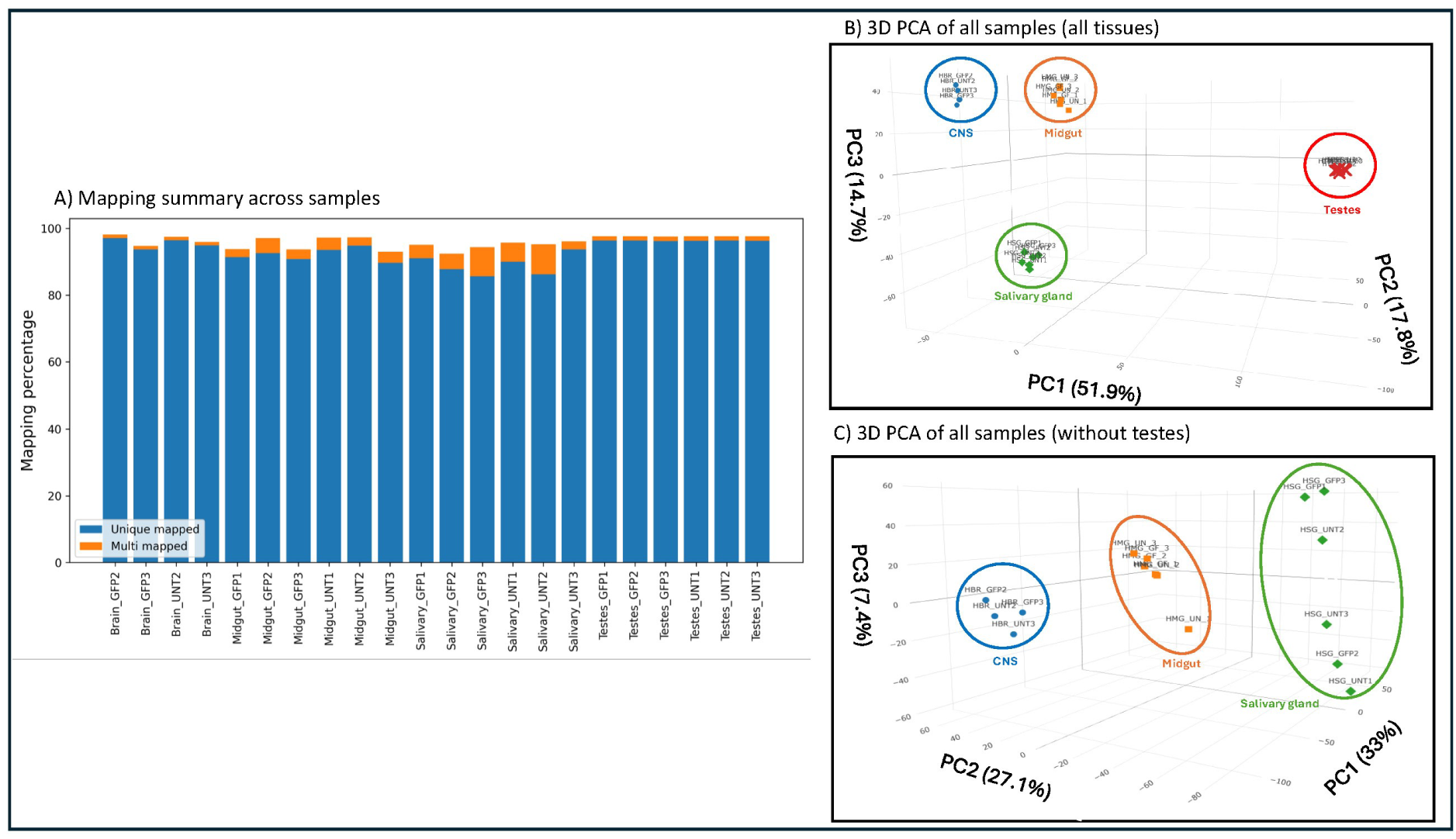
Sequencing quality metrics, sample clustering, and differential expression overview. (A) Mapping efficiency of trimmed, quality-filtered reads across all RNA-seq libraries, showing consistently high alignment rates (median ≥ ∼90%) across tissues and treatment conditions. (B) Three-dimensional principal component analysis (PCA) plot based on variance-stabilised gene expression counts, demonstrating clear clustering of samples by tissue type, with strong separation along the first three principal components. (C) PCA plot excluding testes samples, improving visualization of clustering patterns among the other three tissues.

Principal component analysis (PCA) of variance-stabilised counts revealed clear separation of samples by tissue along the first three principal components, with tight clustering of biological replicates within each tissue group. This pattern confirms both the robustness of tissue dissection and the reproducibility of library preparation (Fig. 2B-C).

### dsGFP injection induces a minimal transcriptional response that is only detectable in a tissue-specific manner

Differential expression analysis showed that dsGFP injection induced a modest transcriptional response. With a relaxed statistical threshold (*padj* <0.05), only 1.64% of genes in the *H. halys* annotated gene set (287 of 17,483 genes) were affected. Of these, 95.6% of unique DEGs were tissue-specific, with only 12 genes shared between tissues (Fig. 3A). The same trend was observed for both upregulated (n = 129) and downregulated (n = 158) genes (Fig. 3B & C; SI_3).

**Figure 3.**
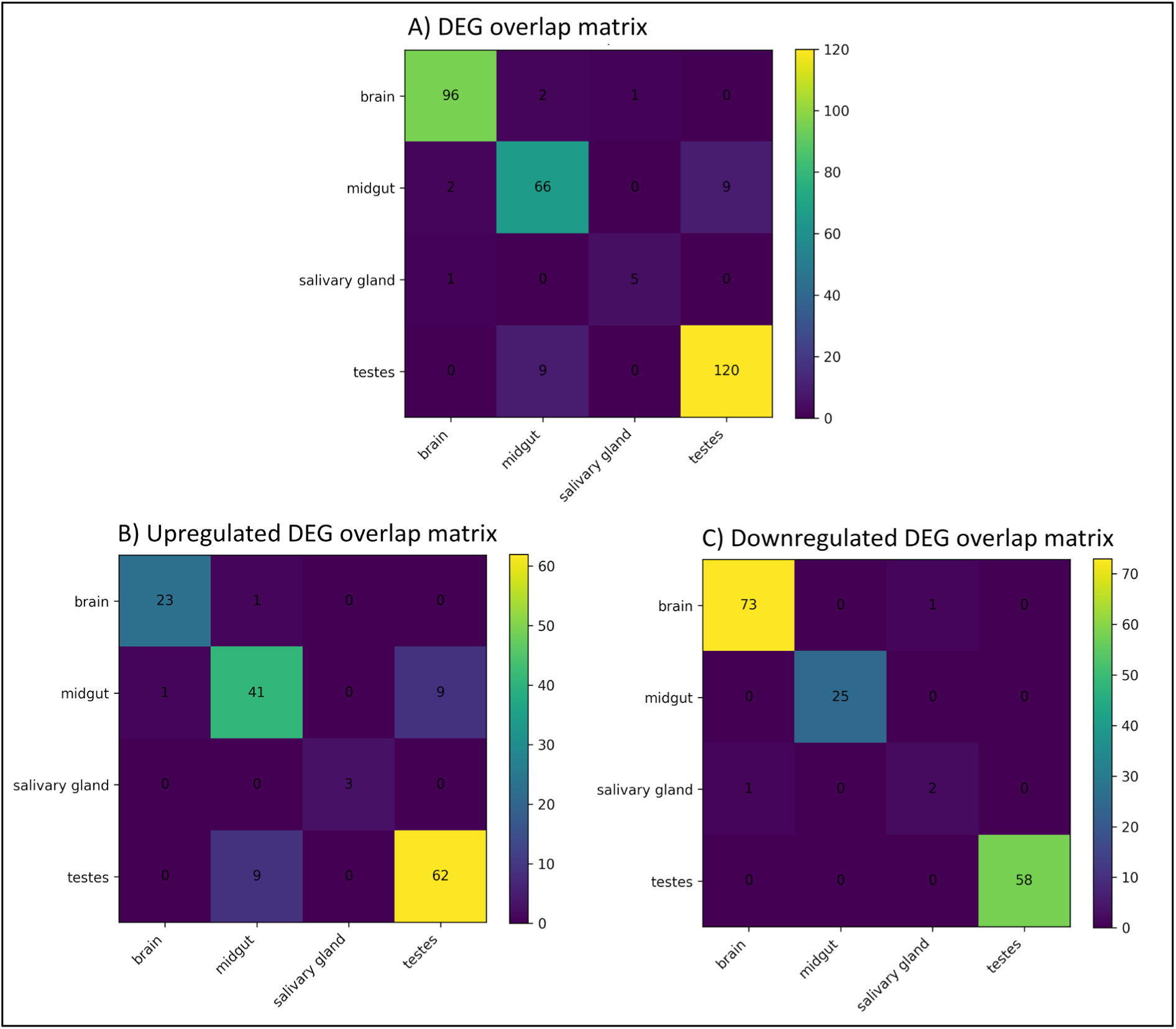
Differentially expressed gene overlap across tissues. (A) Gene overlap matrix showing shared differentially expressed genes (DEGs; p_adj_ <0.05) across tissues in response to dsGFP treatment (GFP vs. untreated control; total n = 287). (B) Overlap matrix of upregulated genes (log₂ fold change > 1; n = 129), indicating predominantly tissue-specific transcriptional activation with limited cross-tissue overlap. (C) Overlap matrix of downregulated genes (log₂ fold change < 0; n = 158), showing minimal shared responses across tissues and reinforcing strong tissue-specific responses.

The DEGs include only two RNAi-related genes: *clathrin heavy chain* and *Apolipoprotein,* which are known to be involved in the uptake of double-stranded RNA by cells [16, 26, 39–41], were upregulated 1.62-fold and 1.82-fold, respectively, in the brain in our dataset. However, with a stricter threshold (*padj* <0.01 and ≥2-fold expression change), which has been informative in validation screening in other insects [42], only 0.49% of genes (85 genes) were affected (SI_3).

Together, these data show that injection of dsGFP against a non-endogenous target sequence induces only a modest transcriptomic effect in wild-caught bugs, consistent with previous transcriptome-scale assessments in other species [28].

### Tissue-resolved expression atlas of core RNAi machinery genes

To characterise the baseline expression landscape of RNAi pathway components, normalised expression values (log₂[mean normalised count + 1], here and in all analyses presented below) from water-injected control libraries were analysed (n = 3 biological replicates per tissue, except n=2 for brain tissue). Candidate genes spanning antiviral, intracellular transport, nuclease, RISC and piRNA, dsRNA uptake, miRNA, and siRNA functional categories were profiled based on previous work [43] (Fig. 4; SI_4).

**Figure 4.**
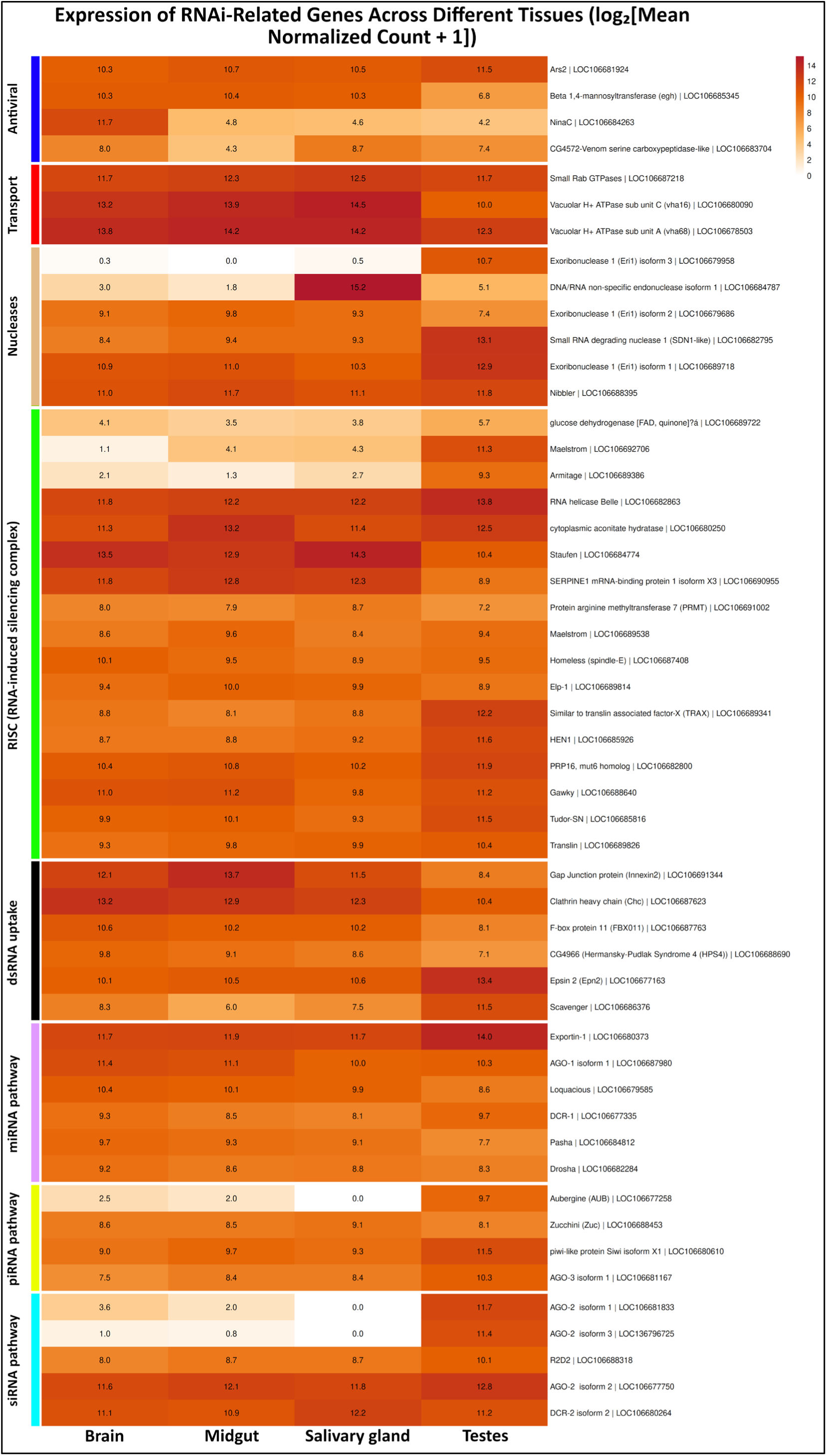
Tissue-resolved expression patterns of RNAi machinery genes. Heatmap showing normalised expression values of genes associated with RNAi pathways across brain, midgut, salivary glands, and testes. Rows represent individual genes grouped by functional categories (antiviral, intracellular transport, nucleases, RISC, uptake, miRNA, piRNA, and siRNA), ordered as displayed in the heatmap, and columns represent tissues. The colour scale indicates expression intensity from low (light) to high (dark). Distinct tissue-specific patterns are observed, including enrichment of RNAi components in testes and strong salivary gland expression of nuclease-related genes.

#### Antiviral genes

Four genes associated with antiviral dsRNA sensing – *Ars2*, *egh* (*Beta 1,4-mannosyltransferase*), *NinaC*, and *CG4572-Venom serine carboxypeptidase-like* – were detected across all four tissues. Expression was broadly distributed, with 3-fold enrichment observed for *NinaC* in brain tissue.

#### Intracellular dsRNA transport

Genes encoding *Small Rab GTPases*, *Vacuolar H⁺-ATPase subunit C*, and *Vacuolar H⁺-ATPase subunit A* were robustly and uniformly expressed across all tissues, suggesting constitutive activity of endosomal acidification and vesicular trafficking machinery, irrespective of tissue identity.

#### Nucleases

Among nuclease-encoding genes, the gene *DNA/RNA non-specific endonuclease* exhibited highly tissue-restricted expression, with strong and exclusive enrichment in salivary glands and near-absent expression in brain, midgut, and testes. This finding aligns with the known *dsRNase* activity previously reported in *H. halys* saliva [14, 44, 45]. In contrast, *Exoribonuclease-1 (Eri1) isoforms 1* and *2* were broadly expressed, though *Eri1 isoform 3* showed strongly elevated expression exclusively in testes, with negligible expression in all other tissues. *Small RNA degrading nuclease 1* (*SDN1-like*) and *Nibbler* were ubiquitously expressed, with moderate enrichment in testes of *SDN1-like (1.5-fold compared to the other tissues)*. Please also see the following results subsection for further details on diverse nuclease gene profiles.

#### RISC-associated and piRNA pathway genes

The majority of genes encoding RISC components, including *RNA helicase Belle, cytoplasmic aconitase hydratase, Staufen, SERPINE1 mRNA-binding protein 1, HEN1, PRP16, Gawky, Tudor-SN,* and *Translin*, were broadly expressed across all four tissues, with overall enrichment in testes for nearly half of these. Furthermore, *Maelstrom*, *Armitage*, and *glucose dehydrogenase[FAD, quinone]-related* transcripts displayed markedly elevated expression in testes with minimal expression in the remaining tissues, suggesting testes-enriched piRNA pathway activity (Fig. 4).

#### dsRNA uptake receptors

Genes encoding known or candidate extracellular and endocytic dsRNA uptake receptors, *Gap Junction protein (Innexin-2), Clathrin heavy chain (Chc), F-box protein 11 (FBX011), Hermansky-Pudiak Syndrome-4 (HPS4), Epsin 2 (Epn2),* and *Scavenger receptor*, were all detectably expressed across the four tissues examined, consistent with broad competence for receptor-mediated dsRNA uptake throughout the body.

#### miRNA pathway genes

The core miRNA biogenesis genes *Exportin-1, AGO-1 isoform 1, Loquacious, DCR-1, Pasha,* and *Drosha* were expressed broadly across all four tissues.

#### siRNA pathway genes and AGO isoform diversity

Key siRNA pathway components, including *AGO-2, R2D2,* and *DCR-2*, were expressed across all tissues. Notably, two additional AGO-2 isoforms *(isoforms 1* and *3*) exhibited testes-specific expression, with no detectable expression in brain, midgut, or salivary glands. Together, these results indicate isoform-specific expression of AGO-2 expression in the testes, which may reflect enhanced siRNA pathway activity in the germline, potentially linked to transposon silencing and genome integrity during spermatogenesis [46].

### Overall patterns of expression of nuclease and protease coding genes

Normalised expression profiles of the specific candidate genes sourced from a previous study [11] were examined for 53 nuclease-encoding and 60 protease-encoding genes spanning exonuclease, endonuclease, ribonuclease, exoribonuclease, nucleotidase, dsRNase, Dicer, trypsin, chymotrypsin, cathepsin, aspartic proteinase, and dipeptidase families (Fig. 5 & 6, Fig. S2, Fig. S3, SI_5, and SI_6).

**Figure 5.**
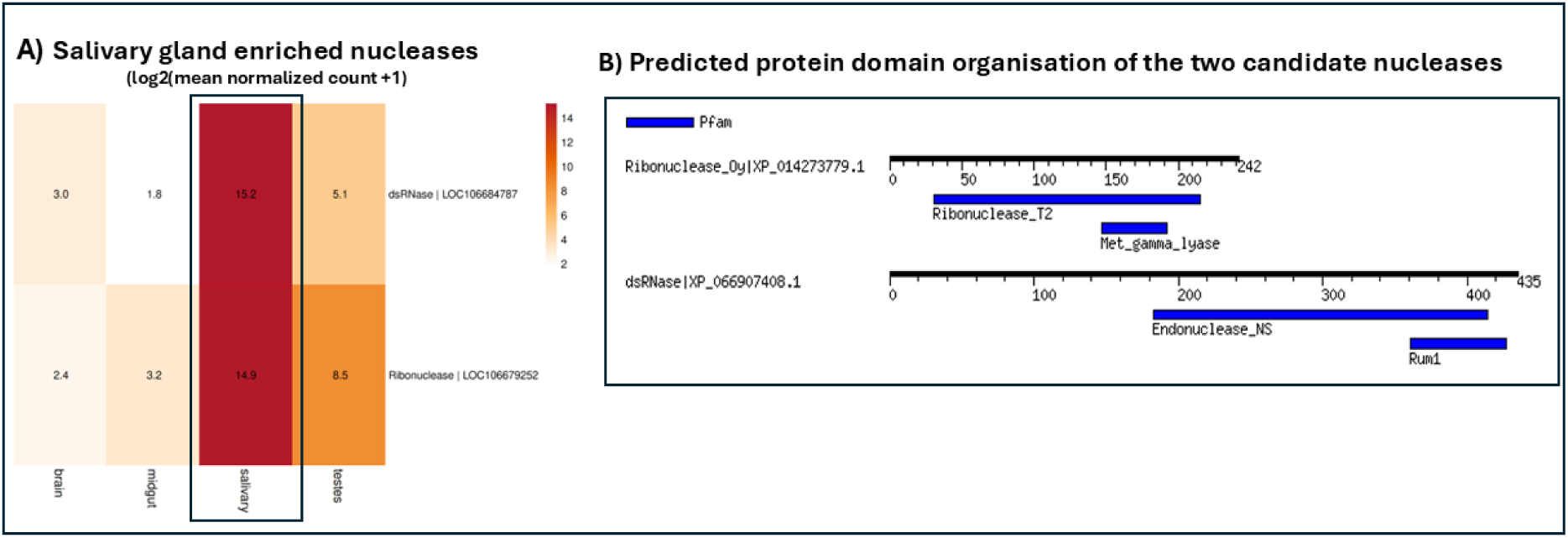
Salivary gland-enriched nuclease candidates and predicted protein domain organisation. (A) Heatmap showing log2 mean normalised read counts (+1) of two candidate nuclease genes across four tissues (brain, midgut, salivary gland, testes). Rows represent genes and columns represent tissues. Numbers within cells indicate expression values. Colour scale ranges from low expression (light) to high expression (dark red). Black outline highlights the salivary gland column. (B) Predicted protein domain organisation of the two candidate nucleases. Horizontal black lines represent full protein length in amino acids, with scale bars shown above each sequence. Blue boxes indicate annotated Pfam domains. Protein names and accession numbers are shown at left. Domains identified include Ribonuclease_T2 in Ribonuclease_Oy (XP_014273779.1), and Endonuclease_NS in dsRNase (XP_066907408.1).

**Figure 6.**
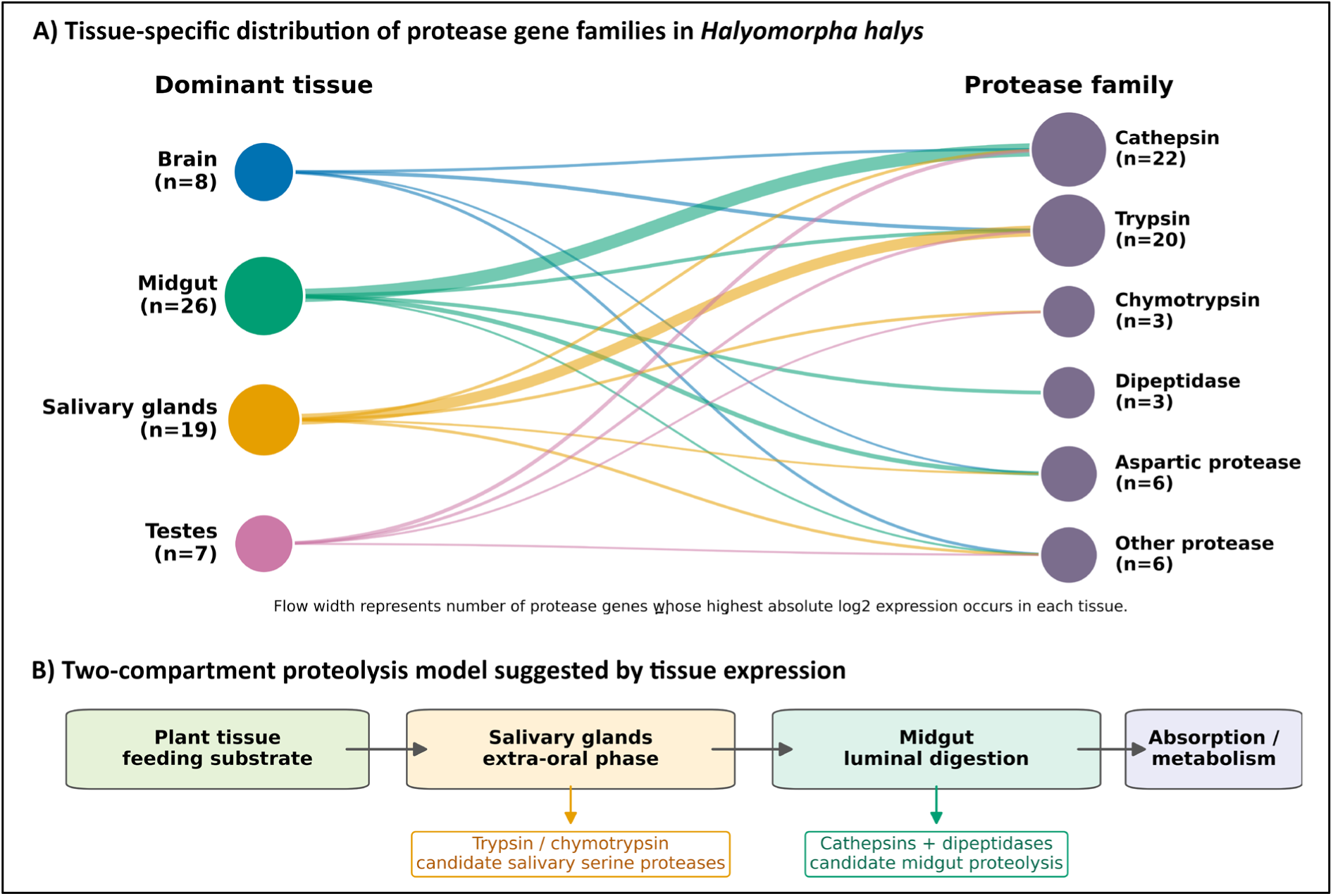
Tissue-specific specialization and inferred digestive roles of protease genes in Halyomorpha halys. (A) Alluvial diagram showing the distribution of protease gene families according to the tissue in which each gene exhibited the highest absolute normalized expression (log2 mean normalized count + 1). Flow width is proportional to the number of genes assigned to each tissue–family combination. Major families included cathepsins, trypsins, chymotrypsins, dipeptidases, aspartic proteases, and other proteases. Midgut-enriched genes were dominated by cathepsins, whereas salivary glands showed strong representation of serine proteases (trypsin/chymotrypsin), suggesting functional partitioning of proteolysis. (B) Conceptual model of a two-compartment digestive proteolysis system inferred from tissue expression profiles. Salivary gland-enriched trypsin/chymotrypsin genes are proposed to participate in extra-oral digestion during feeding, whereas midgut-enriched cathepsins and dipeptidases likely mediate luminal digestion and downstream nutrient processing. Dominant tissue assignments were based on highest absolute expression values. Tissue-enriched candidates were identified using row z-score ≥ 1.0 together with absolute log2 expression ≥ 3.0.

Genes exhibited diverse expression patterns across the nuclease family (Fig. S3). *Nucleotidase | LOC106682044* was the sole gene with high expression across all tissues (13- to 14-fold enrichment), suggesting it plays a constitutive role independent of tissue context. In contrast, many nucleases have peak expression in the testes, being either testes-exclusive (*Exonuclease | LOC106683836* and *Exoribonuclease | LOC106679958*) or being expressed across all tissues but with highest levels in the testes (∼13-fold enrichment), indicating elevated but not exclusive activity in this tissue. Only a small number of nucleases showed higher expression in the brain relative to other tissues, and these were expressed at moderate levels (∼9-fold enrichment), indicating that brain-associated nuclease activity is comparatively limited in both magnitude and gene representation.

Intriguingly, two nucleases exhibited exclusive enrichment in the salivary glands: *dsRNase* and *Ribonuclease-Oy-like* (Fig. 5A, ∼15-fold enrichment). This restricted expression pattern supports a potential specialised role in salivary gland function. Going further, functional characterisation of these salivary-enriched candidates revealed conserved catalytic domains, consistent with nuclease activity (Fig. 5B, Fig. S2). However, signal peptide predictions suggest distinct sites of activity for these proteins.

The dsRNase contains a high-confidence signal peptide with a predicted cleavage site between positions 20–21 (Fig. S2D), consistent with secretion via the classical pathway. In contrast, Ribonuclease-Oy-like lacks a detectable signal peptide (Fig. S2C), indicating that it is unlikely to follow the same secretion route and may function intracellularly or via an alternative mechanism.

Overall, the data resolve three major expression patterns within the nuclease repertoire: (i) strongly testes-enriched genes with either exclusive or predominant expression, (ii) salivary gland-specific nucleases with almost-exclusive expression, and (iii) genes with relatively uniform abundance across tissues. A complete overview of expression profiles for all nuclease genes is provided in the supplementary material (Fig. S3, SI_5).

### Protease-coding genes display tissue-specialized expression consistent with functional compartmentalization of digestion

Protease gene expression was predominantly tissue-specific, particularly in the salivary gland and midgut as sites of digestive enzyme activity. Only a handful of proteases showed relatively uniform expression across tissues, including a subset with highest expression in the brain; these likely have conserved roles in general protein turnover rather than digestive functions [47]. Testes protease expression was generally low and lacked a distinctive gene subset.

In contrast, five trypsin paralogs and two chymotrypsins were strongly and exclusively enriched in the salivary glands (∼14–17-fold enrichment; Fig. 6A), consistent with extracellular proteolytic activity during feeding. This restricted expression pattern aligns with previous observations and indicates a specialised salivary serine protease complement [11].

The midgut exhibited a distinct expansion of lysosomal and digestive proteases. Eleven cathepsin L1 and L1-like paralogs showed strong midgut enrichment (∼13–16 fold), with minimal expression in the brain or salivary glands. Additional midgut-dominant proteases included cathepsin D (LOC106690635), cathepsin B-like (LOC106689147), and two aspartic proteinases (LOC106689815, LOC106679181). The consistently high expression of these genes supports a concentrated proteolytic environment associated with intracellular digestion [48].

The protease repertoire resolves into four principal expression patterns: (i) salivary gland-restricted serine proteases, (ii) midgut-enriched lysosomal proteases, (iii) broadly expressed housekeeping proteases, and (iv) a limited set of brain- or testes-associated genes. This organisation reflects functional compartmentalization of proteolysis across tissues (Fig. 6A-B). Full expression matrices and individual gene values are provided in SI_6 and Fig. S4.

### Tissue-specific expression modules among detoxification gene families

To characterize tissue-specialized functions of major detoxification enzyme superfamilies in *H. halys*, normalized expression profiles for 148 detoxification-associated genes were analysed (Fig. 7a-c, Fig. S6, SI_7). Specifically, the CYP superfamily represented the largest detoxification cohort (71 genes, 48.0% of total), dominated by CYP3-clan members (37 genes) followed by CYP4-clan genes (25 genes), with smaller CYP2-clan (6 genes) and mitochondrial clan (MIT, 3 genes) contingents. GSTs comprised 26 genes (17.6%). Esterases represented the largest family by diversity (51 genes, 34.5%), subdivided into Esterase FE4-like (22 genes), venom carboxylesterase-6-like (19 genes), Esterase E4-like (5 genes), and minor carboxylesterase variants. Detailed gene counts and tissue-specific expression values for all 148 genes are provided in SI_7 and Fig. S5.

**Figure 7.**
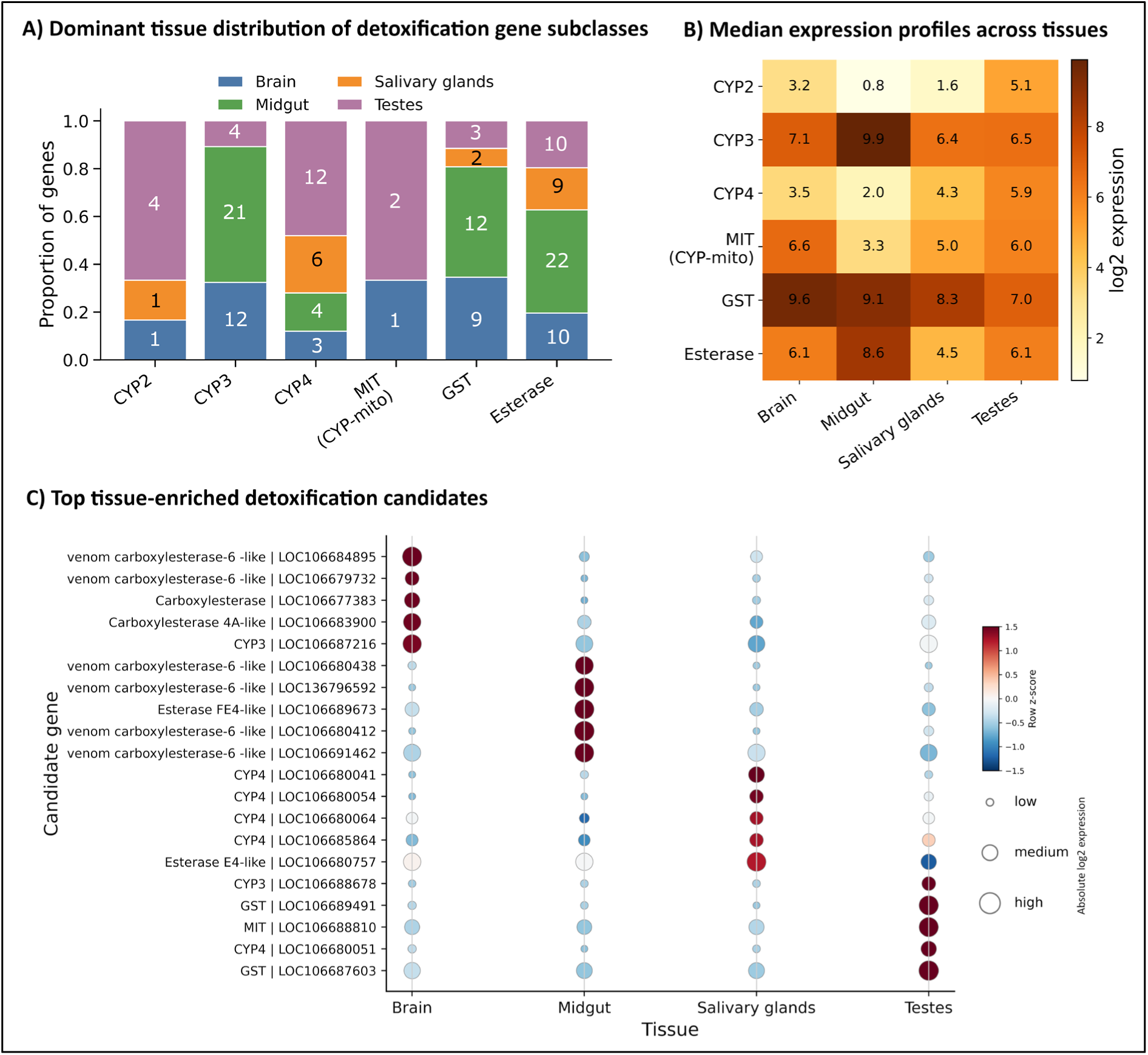
Tissue-specific organization of detoxification-associated gene subclasses in Halyomorpha halys. Transcript abundance profiles of detoxification-associated genes were summarized across brain, midgut, salivary glands and testes to identify subclass- and tissue-level expression patterns. (A) Proportion of genes in each detoxification subclass whose highest absolute log₂-normalized expression occurred in each tissue, revealing dominant tissue distributions among CYP2 (n=6), CYP3 (n=37), CYP4 (n=25), mitochondrial CYPs (n=3), GSTs (n=26) and esterases (n=51). (B) Heatmap of median absolute log₂-normalized expression values for each subclass across tissues, showing broad subclass-level expression patterns. (C) Bubble plot of the top five tissue-enriched candidate genes per tissue, selected using row-wise z-score-transformed expression values. Bubble color indicates relative enrichment across tissues for each gene, and bubble size indicates absolute log₂ expression. Genes were classified into detoxification subclasses based on annotation names. Dominant tissue assignment and subclass medians were calculated from the absolute log₂-normalized expression matrix, whereas tissue-enriched candidates were selected from the row-wise z-score matrix.

#### Cytochrome P450 monooxygenases

For the large CYP superfamily (SI_7), hierarchical clustering resolved distinct tissue-associated expression modules for the different subfamily clans. CYP3 genes were predominantly assigned to the midgut-dominant category, whereas CYP2 and mitochondrial CYP genes showed stronger testes representation (Fig. 7A). Median expression profiles further showed that CYP3 had its highest subclass-level expression in the midgut, while CYP2 expression was highest in testes (Fig. 7B).

A subset of CYP3-clan genes showed high brain expression, including strongly brain-biased genes such as LOC106687998, LOC106691459, LOC106680992 and LOC106680994 (Fig. S5; SI_7). Additional CYP3 paralogs, including LOC106686595, LOC106690280 and LOC106690578, were highly expressed in brain but also retained substantial expression in midgut and/or salivary glands, indicating broader tissue activity rather than strict brain specificity (SI_7). Similar nervous system-associated expression of cytochrome P450 genes has been reported in other insects, where certain CYPs are proposed to contribute to local detoxification and neuroprotection within neural tissues [49, 50].

A larger midgut-enriched CYP module was also evident, including 18 CYP3 genes with log₂ midgut expression >10 (Fig. S5; SI_7). Highly expressed representatives included CYP3 | LOC106690077, CYP4 | LOC106678588 and CYP3 | LOC106678897 (SI_7). This midgut-biased CYP expression is consistent with a role in digestive xenobiotic metabolism, as insect CYP3 and CYP4 clans are commonly enriched in lineage-specific paralog expansions associated with xenobiotic metabolism and pesticide response [51] (Fig. 7A,B).

Several CYP2&4 genes showed testes-biased expression (Fig. 7A, C; SI_7). CYP4 | LOC106677250 was strongly enriched in testes but retained moderate midgut expression, whereas CYP2 | LOC106686814, CYP4 | LOC106679207 and CYP4 | LOC106680051 showed low or undetectable midgut expression and higher testes expression (SI_7). This pattern suggests that a subset of CYP2 and CYP4 genes may contribute to reproductive or endogenous metabolic processes. Such an interpretation is consistent with the broader observation that CYP2 and mitochondrial CYP genes are often more conserved than the CYP3/CYP4 detoxification clans that often exhibit lineage-specific expansions [51]. Salivary gland-specific CYP expression was limited, with CYP4 | LOC106680041 showing the clearest salivary-enriched profile among CYPs (Fig. 7C; SI_7).

Of the three mitochondrial clan CYP genes that were detected (SI_7), LOC106688810 showed pronounced testes enrichment and was among the top testes-enriched detoxification candidates (Fig. 7C), with log₂ expression of 12.9 in testes compared with 7.2 in brain, 6.8 in midgut and 7.3 in salivary glands (SI_7). The remaining two mitochondrial CYP paralogs, LOC106678426 and LOC106689460, were weakly and broadly expressed across tissues (SI_7).

#### Glutathione S-transferases

A total of 26 GST genes were detected and were broadly distributed across the four tissues (SI_7), with high general expression compared to other detoxification gene families (CYP clans and esterases). Proportionally, GST dominant-expression assignments were mainly distributed between brain and midgut, with a smaller salivary component (Fig. 7A-B). LOC106688542 and LOC106685622 were the most highly and constitutively expressed GSTs, reaching log₂ expression values of 14.9 and 12.4, respectively (SI_7). Tissue-restricted expression was most pronounced for LOC106689491, which was among the top testes-enriched detoxification candidates (Fig. 7C; SI_7). Brain-enriched GSTs included LOC106680062 and LOC106679352, while LOC106688241 and LOC106687603 showed highest expression in midgut and testes, respectively (SI_7). Overall, the broad expression of GSTs suggests a constitutive conjugation and antioxidant defence capacity across tissues, particularly in the brain and midgut (Fig. 7A,B). This interpretation is consistent with established roles of insect GSTs in xenobiotic conjugation, insecticide response and protection against oxidative stress [52, 53].

#### Carboxyl/hormone esterases

In total, 51 esterase-associated genes were identified (SI_7). Expression patterns were dominated by a large midgut-enriched cohort (Fig. 7A-B). Approximately 20 genes showed peak midgut expression above log₂ 10 (Fig. S5; SI_7). Highly expressed representatives included Esterase FE4-like | LOC106689685 and several venom carboxylesterase-6-like paralogs, including LOC106680412, LOC136796592, LOC106680438 and LOC106680420 (Fig. 7C; SI_7). This midgut expression pattern is consistent with candidate roles in hydrolytic metabolism of dietary or xenobiotic substrates (Fig. 7A–C). Insect carboxyl/cholinesterases are multifunctional enzymes involved in hydrolytic biotransformation, pesticide resistance, host-plant adaptation and chemical ecology [54, 55].

Among the other three tissues, salivary gland-enriched esterases included Esterase E4-like | LOC106680757 and venom carboxylesterase-6-like | LOC106692443, although LOC106680757 also retained high expression in brain and midgut (Fig. 7C; SI_7). Given the importance of saliva in *H. halys* feeding injury and plant interaction, these salivary-enriched esterases may represent candidate enzymes involved at the plant–insect feeding interface [56, 57]. In the testes, Juvenile hormone esterase-like | LOC136797101 was the dominant esterase isoform, accompanied by a near-exclusively testes-expressed venom carboxylesterase-6-like gene, LOC106692607 (Fig. 7C; SI_7). The testes enrichment of juvenile hormone esterase-like expression suggests a possible role in local hormone-associated metabolism, although functional validation would be required to confirm juvenile hormone catabolism in this tissue. Lastly, venom carboxylesterase-6-like | LOC106684895 was identified among the top brain-enriched detoxification candidates (Fig. 7C; SI_7).

In sum, detoxification-associated genes in *H. halys* are not uniformly expressed across tissues but are organized into tissue-specific metabolic modules. The midgut appears to be the dominant site of digestive xenobiotic metabolism, combining CYP3-mediated oxidation, GST-mediated conjugation and esterase-mediated hydrolysis. The brain maintains a strong antioxidant/conjugation signature; salivary gland specific genes are likely relevant to host plant interaction; and the testes exhibit a distinct CYP2/CYP4/mitochondrial CYP-dominated expression profile that may be associated less with xenobiotic detoxification and more with endogenous reproductive metabolism, including hormone and lipid metabolism, sperm development, and other physiological processes required for reproduction.

### Tissue-specific expression patterns of aminopeptidase and carboxypeptidase genes

Aminopeptidases and carboxypeptidases are key exopeptidases that facilitate the completion of protein digestion through the sequential removal of terminal amino acids. In insects, these enzymes also play a role in a variety of physiological processes, including nutrient assimilation, metabolism, and reproduction [58]. Normalised expression profiles were examined for 17 aminopeptidase-encoding and 16 carboxypeptidase-encoding genes (Fig. 8a-b).

**Figure 8.**
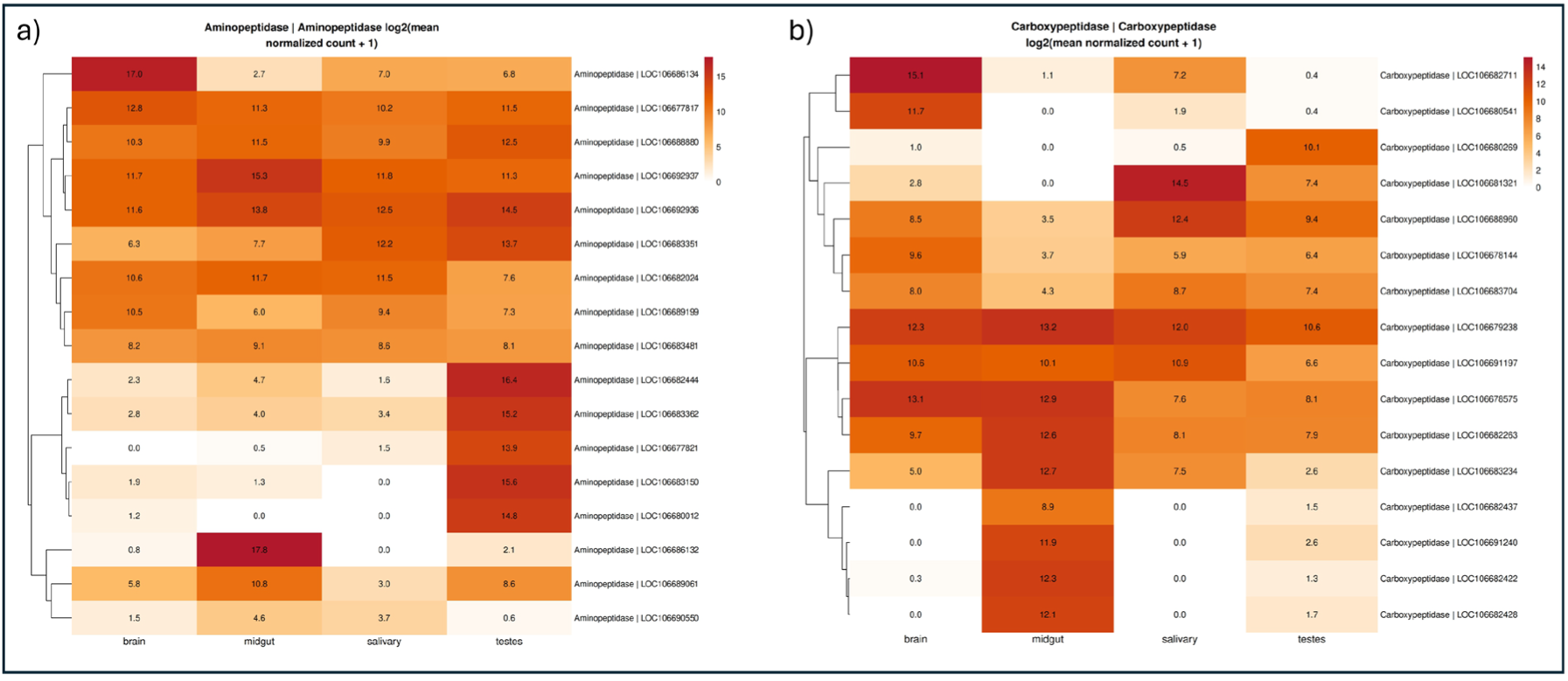
Heatmaps showing normalised expression values of digestive enzyme genes across tissues. (A) Aminopeptidases. (B) Carboxypeptidases. Rows represent individual genes clustered by expression similarity, and columns represent tissues. The colour scale indicates expression intensity from low (light) to high (dark).

The 17 aminopeptidase-encoding genes vary substantially for the extent to which they are generally strongly expressed and with particularly high levels in a given tissue or whether expression was largely exclusive to a single tissue. Nearly half of the aminopeptidases (8 of 17) show peak expression in the testes, with five of these being testes-exclusive while others also had notable salivary gland co-expression or were generally expressed at high levels across all four tissues. Along similar lines, three aminopeptidases had peak expression in the brain and six in the midgut. Notably, no aminopeptidase showed salivary gland-dominant expression, whereas three aminopeptidases showed broad constitutive expression (enrichment score >9 in all tissues; Fig. 8A).

The carboxypeptidase repertoire exhibited clear tissue-partitioned expression patterns, with most genes showing enrichment in either the midgut, brain, or salivary gland (Fig. 8B). Midgut-enriched expression was the most common pattern, observed in seven genes, whereas four genes showed peak expression in the salivary gland and four in the brain. In contrast, only one gene exhibited testes-dominant expression. Several carboxypeptidases displayed strong tissue-specificity, while others showed co-expression across two or more tissues. Notably, LOC106679238 was the only broadly and highly expressed gene across all tissues (brain 12.32, midgut 13.17, salivary gland 11.99, testes 10.58), suggesting a conserved housekeeping or general digestive role.

### Neuropeptide precursor and receptor genes: nuanced expression in the brain and beyond

To characterise tissue-specific neuroendocrine signalling potential, normalised expression profiles were analysed for 33 neuropeptide precursor genes and 49 GPCR genes representing biogenic amine, neuropeptide, LGR/glycoprotein hormone, opsin/photoreceptor, purinergic, adhesion, and orphan receptor families (Fig. 9, Fig. S6, SI_8, and SI_9).

**Figure 9.**
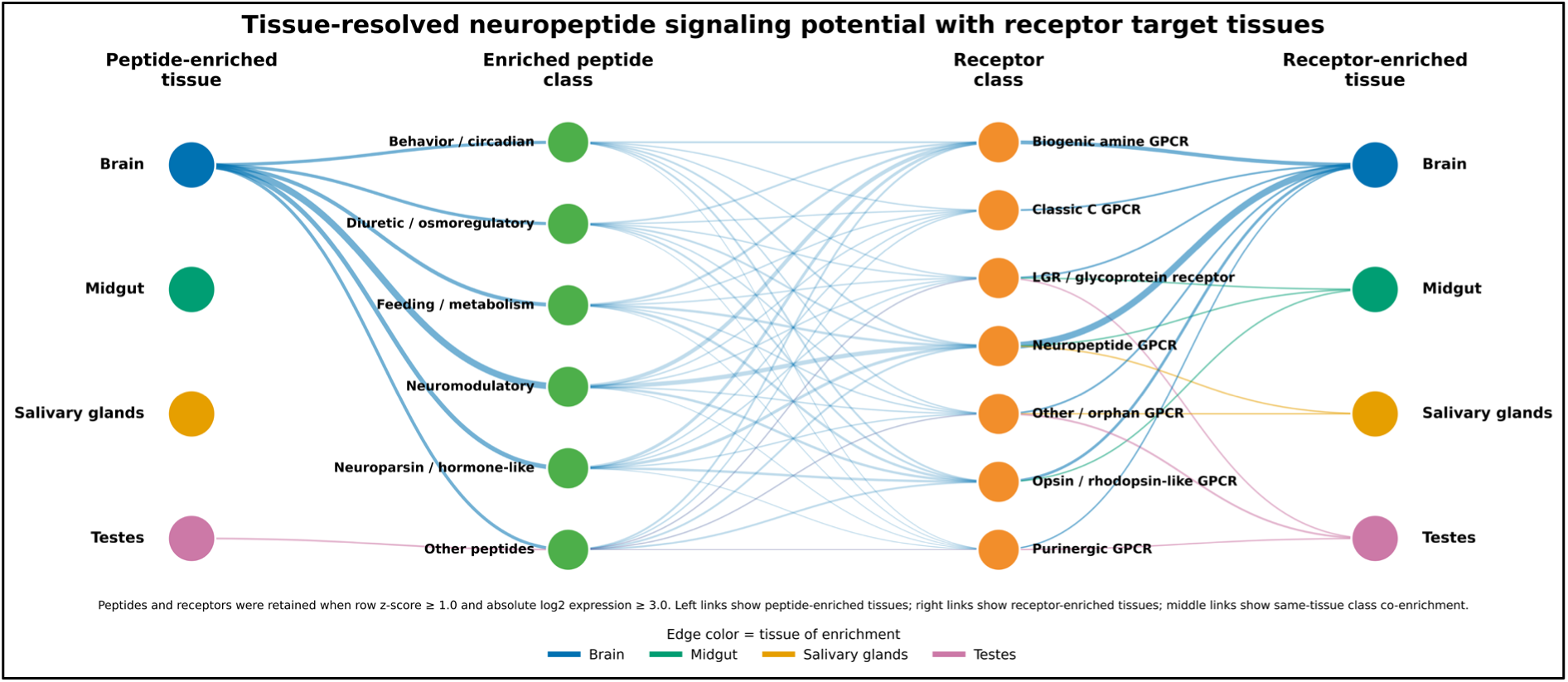
Tissue-resolved neuropeptide signaling potential with receptor target tissues in Halyomorpha halys. Four-column network summarizing tissue-specific enrichment of neuropeptide ligands and receptors across brain, midgut, salivary glands, and testes. Left column indicates tissues enriched for neuropeptide genes (“peptide-enriched tissue”), middle-left column shows functional classes of enriched neuropeptides, middle-right column shows receptor classes, and right column indicates tissues enriched for receptor genes (“receptor-enriched tissue”). Nodes represent tissues or gene classes; node positions are organized by biological category. Colored edges denote the tissue in which enrichment was detected (brain, blue; midgut, green; salivary glands, orange; testes, magenta). Edge width is proportional to the number of genes supporting each connection. Peptide and receptor genes were considered enriched when row-wise z-score ≥ 1.0 and absolute expression ≥ log2(normalized count + 1) of 3.0. Middle connections represent class-level co-enrichment within the same tissue and indicate putative signaling routes rather than experimentally validated ligand–receptor binding. The network highlights the brain as the dominant neuroendocrine signaling hub, with additional tissue-specific signaling modules in testes and more limited receptor-associated modules in midgut and salivary glands.

#### Neuropeptide precursor genes

As expected, expression of neuropeptide precursors was strongly biased toward the brain, with most genes either exclusively detected or markedly enriched in this tissue. Eight precursors, *Proctolin*, two *Neuroparsin* paralogs (*LOC106682902*, *LOC106682903*), *SIF-amide*, *CCH-amide 1, Corazonin, Adipokinetic hormone*, and *RYamide*, were detected only in the brain, indicating strict centralised production.

Several of the highest expressed precursors also showed evidence of multi-tissue expression rather than strict neural restriction. Both *NVP-like* (brain 15.56, midgut 10.47) and *Orcokinin* (brain 14.87, midgut 10.87) displayed substantial expression in both brain and midgut. The comparable magnitude of expression across these tissues may support a dual role in central signalling and peripheral (likely enteroendocrine) regulation.

A subset of precursors showed broad distribution across multiple tissues. *CRF-like Diuretic Hormone A* (brain 11.99, midgut 9.37, testes 8.79) and *Myosuppressin* (brain 11.45, midgut 10.39, testes 7.75) were consistently expressed at high levels in three tissues except salivary gland tissue, indicating systemic signalling roles rather than tissue-restricted functions.

Salivary gland expression was evident for several neuropeptides, although in most cases this occurred alongside comparable or higher brain expression. *IDLSRF-like* (salivary 11.95, brain 12.10*), Insulin-like peptide* (salivary 9.52, brain 8.82), *CNM-amide* (salivary 8.63, brain 9.20), *Tachykinins* (salivary 8.36, brain 12.53), and *Ion Transport peptide* (salivary 7.53, brain 8.65) all showed substantial salivary expression, supporting an active role for peptidergic signalling in this tissue, consistent with its secretory function during feeding.

*Ecdysis triggering hormone* was the only precursor with peak, albeit moderate, expression in testes (5.79), while remaining low in other tissues (brain 2.14, midgut 1.56, salivary 1.99). This restricted pattern suggests a specialised reproductive or developmental role distinct from its canonical function in moulting [59].

#### Neuropeptide receptor genes

Analysis of receptor expression similarly revealed strong tissue specificity, with the brain representing the principal site of GPCR expression. Twenty-one receptors showed clear brain-dominant profiles (log₂ brain > 7 with minimal expression in other tissues), spanning multiple receptor classes.

CNS-restricted expression encompasses all biogenic amine receptors, including *dopamine* (DR2, DR4, DR5), *octopamine* (OA-1, OA-2), *tyramine*, and *serotonin receptors*, were highly expressed in the brain (log₂ 7.95–10.89) with negligible or undetectable expression in midgut and salivary glands. Equally, opsin receptors (*Opsin-1, Opsin-2, Opsin-3*) were exclusively or strongly enriched in the brain, with no detectable expression in midgut or salivary glands, consistent with photoreceptive signalling capacity being restricted to neural tissues.

For some receptor families, different paralogs were associated with different tissues. For example, *GPCR moody* genes were either strongly expressed in the salivary glands (*LOC106688042*; 10.65) or in the testes (*LOC106690378*; 8.77). Similarly, the LGR family exhibited clear functional partitioning across tissues. The *glycoprotein hormone receptor* (*LOC106686122*) was highest in testes (12.07), suggesting a reproductive role, whereas the *FSH-like receptor* (*LOC106685828*) was predominantly expressed in midgut (10.12), and *Methuselah* showed co-enrichment in midgut (9.87) and testes (9.65), indicating tissue-specific diversification within this receptor class.

Additional receptors also showed enrichment in the salivary glands, including *FMRFamide receptor* (*LOC106685288*; 9.12), and *Trissin receptor* (*LOC106686894*; 8.39). Although also expressed in the brain, their elevated salivary expression supports a functional role for GPCR-mediated peptidergic signalling in this tissue.

Further testes-enriched receptors included the *glycoprotein hormone LGR* (12.07), *Adenosine receptor-1* (7.68), and *Ecdysis-triggering hormone receptor-1* (7.65). These patterns indicate that reproductive tissues possess a distinct complement of neuropeptide and hormone receptors, consistent with active local signalling networks.

In contrast, the adhesion GPCR *Latrophilin* (*LOC106684418*) showed consistently high expression across all tissues (brain 11.79, midgut 10.29, salivary 11.59, testes 7.11), distinguishing it as a broadly utilised receptor likely involved in general cellular or synaptic functions rather than tissue-specific signalling.

#### Specific neuropeptide signalling capacities in target tissues

Integration of neuropeptide precursor and receptor genes enriched in specific tissues (row-wise z-score ≥ 1.0; log₂(normalized count + 1) ≥ 3.0) revealed tissue-structured signalling network connectivity (Fig. 9, SI_11).

The brain emerged as the primary neuroendocrine hub, exhibiting the highest diversity of enriched peptide classes, receptor classes, and putative signalling interactions. Brain-enriched modules encompassed behaviour/circadian, neuromodulatory, feeding/metabolic, and diuretic–osmoregulatory peptides, predominantly linking to biogenic amine, neuropeptide, classic C, and LGR/glycoprotein GPCR classes (Fig. 9).

Peripheral tissues displayed smaller, specialized modules. The midgut showed enrichment primarily in receptor-mediated links, aligning with its role in peptide-responsive digestion and metabolic regulation. Salivary glands featured a restricted module involving feeding-related peptides and neuropeptide/orphan GPCRs, consistent with local secretory control. Testes formed a distinct reproductive module enriched for select peptides and receptors, including purinergic, opsin/rhodopsin-like, and orphan GPCRs.

This network underscores centralized neuropeptide signalling in the brain, with peripheral tissues maintaining few selective receptor repertoires and limited ligand synthesis for specialized functions.

### Distinctive classes of gene enrichment in each of the salivary glands, midgut, brain, and testes

To quantify gene expression specificity across the four profiled tissues, the Tau (τ) metric was applied to CPM-normalized expression data from the control tissue samples (see Methods for statistical thresholds). A total of 4,072 genes were classified as strictly tissue-specific, with the testes exhibiting the largest complement (n=2,126 genes; 52.2%), followed by the brain (n=887 genes; 21.8%), midgut (n=610 genes; 15.0%), and salivary glands (n=449 genes; 11.0%) (Fig. 10, the list of tissue specific gene-subsets are provided in supplementary file SI_12). The disproportionately large testes-exclusive transcriptome is consistent with the extensive spermatogenic-specific gene expression programs reported across insects and other metazoans [60, 61]. The 20 highest-expressed tissue-specific genes per tissue are summarized in Table S1 in Supplementary File SI_1, with the full gene list in the supplementary file SI_12. Then, to assess the tissue-specific gene sets globally, gene ontology (GO) enrichment analysis was performed (Fig. 10, Figs. S8–S11 in SI_1 file).

**Figure 10.**
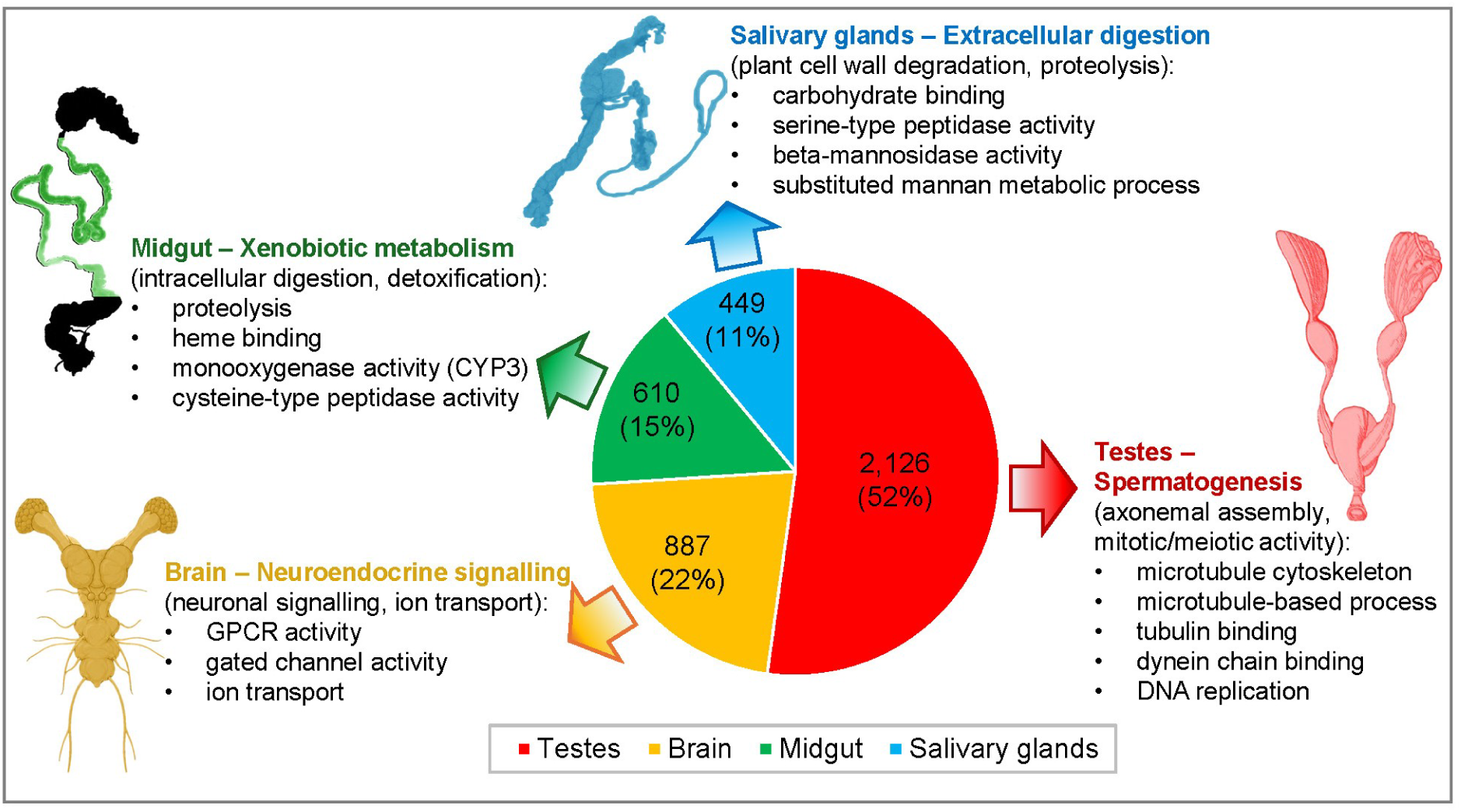
Summary of tissue-specific gene expression in H. halys adult males. The fraction of all tissue-specific genes attributed to each tissue is indicated (gene count and percentage), with key GO enrichment terms and overall biological features of each tissue indicated. Tissue schematics are based on Fig. 1 and [62].

Salivary gland tissue-specific enrichment showed the most focused functional profile of all tissues, dominated by beta-mannosidase activity, serine-type peptidase activity, and carbohydrate binding (Fig. 10, and Fig. S7). This GO signature reflects the specialized role of hemipteran saliva in enzymatic degradation of plant cell walls during feeding and host-plant penetration, as part of an overall functional requirement for carbohydrate hydrolysis and extracellular proteolysis.

The midgut-specific gene set (for the M2-M3 gut regions, Fig. 1A) was enriched for proteolytic activity and xenobiotic metabolism, reflecting the midgut’s established roles in intracellular digestion and aligning with the pronounced midgut enrichment of the CYP3-clan cytochrome P450 genes and cathepsin proteases, as identified in the detoxification and protease profiling sections (Fig. 7, Fig. S8).

The GO enrichment signature for the brain tissue samples, centred on GPCR signalling and ion channel activity, is consistent with the complex neuroendocrine signalling functions of the insect brain and aligns with the brain-enriched neuropeptide receptor and biogenic amine signalling networks identified in the neuropeptide profiling section (Fig. 9, Fig. S9).

Finally, the testes yielded the highest number of tissue-specific genes, with the broadest enrichment profile, spanning from ion binding and kinase activity to multiple terms associated with DNA replication and cytoskeletal regulation and dynamic activity (Fig. S10). This GO enrichment signature likely reflects the extensive axonemal assembly, mitotic, and meiotic activity characteristic of the active spermatogenic program in *H. halys* males.

## Discussion

In parallel with its growing invasive pest status, genomic and transcriptomic resources for *H. halys* have expanded substantially, with the 1.15-Gb genome assembly, stage- and sex-resolved whole-body transcriptomes, and organ-focused salivary gland and gut datasets [6, 10, 11, 36]. Within this landscape, the present work provides a tissue-resolved, multi-organ RNA-seq atlas for adult males spanning brain, midgut (M2-M3 region), salivary glands, and testes, explicitly structured around RNAi machinery, nucleases, detoxification, digestion, and neuroendocrine signalling. This positions *H. halys* alongside model and non-model insects where tissue-level or cell-type–level atlases now serve as primary references for target discovery and functional investigations [30, 63, 64].

### Relation to existing H. halys genomic and transcriptomic resources

Sparks et al. (2020) and others used the *H. halys* genome to highlight expansions in chemoreceptors, cathepsins, and detoxification genes as putative determinants of polyphagy and insecticide resistance [6, 11, 18, 22]. The present atlas refines those genome-level inferences by assigning many of these gene families to specific tissues. GPCR signalling mechanisms, neuropeptides, and several odorant binding proteins are strongly brain-enriched. Meanwhile, cathepsins and carboxyl/hormone esterases are predominantly midgut- or testes-biased, and chemosensory and visual genes (OBPs, opsins) are almost entirely restricted to the brain. Similarly, the developmental and immune-induced transcriptomes in Sparks et al. (2014) catalogued whole-body developmental RNA-seq to identify putative core innate immunity and RNAi pathway components [10]. Going further, our data demonstrate that virtually all major siRNA, miRNA, and piwiRNA factors (DCR-1/2, AGO-1/2, R2D2, Drosha, Pasha) are constitutively expressed across all four adult tissues, with an additional layer of testes-specific diversification of *Argonaute-2* isoforms and few key piwiRNA factors. A similar pattern, with piwiRNA pathway enrichment in the testes in response to viral infection, was previously reported in the mosquito *Aedes albopictus*, indicating a strong antiviral role for the piwiRNA pathway in the reproductive organs [65]. Together, these results move from “presence/absence” of candidate pathways in previous resources towards a spatially explicit map of where these pathways are likely to be most active in adult males.

### RNAi competence, dsRNase barrier, and our data in the context of prior H. halys RNAi studies

Several studies have established that *H. halys* is highly RNAi-competent when dsRNA is injected directly into the haemolymph. Lu et al. (2017) showed robust embryonic and maternal RNAi by targeting *Sex combs reduced* [13], and Mogilicherla et al. (2018) identified multiple lethal adult targets by injection, confirming that systemic dsRNA can generate strong pest-control–relevant phenotypes [15]. Recent formulation work further demonstrated that dsRNA against *Clathrin heavy chain* is almost fully lethal by injection, whereas the same construct is ineffective orally unless protected from salivary nucleases with dsDNA, highlighting non-specific endonuclease-mediated dsRNase activity as the dominant barrier to oral RNAi in this species [14].

Against this background, the present dataset shows that haemolymph-injected non-specific dsRNA (dsGFP) induces only minimal transcriptomic changes and does not upregulate canonical RNAi core genes in any of the four tissues examined; the only consistent signal within the pathway is a subtle increase in *clathrin heavy chain* expression in brain. This contrasts with species such as *Acyrthosiphon pisum* and *Manduca sexta*, where exogenous dsRNA clearly induces *dcr2/ago2* and other RNAi components, and with work in *Schistocerca gregaria* showing that tissue differences in *dicer-2/argonaute-2* expression correlate with systemic RNAi sensitivity [23, 24, 66]. A recent multi-species analysis further emphasises that dsRNA uptake is strongly tissue dependent, even within a single insect [26].

The modest response in our current study in *H. halys* may partly reflect biological heterogeneity of field-collected adults (variation in age, prior infections, or chronic exposure to environmental dsRNA), which could have pre-elevated or desensitised RNAi pathways in some individuals, thereby blunting additional dsGFP-induced transcriptional changes. Nevertheless, taken together with previous injection and formulation studies [28], our results support a model in which *H. halys* relies predominantly on constitutive RNAi machinery, while RNAi resistance during oral-delivery is driven mainly by extracellular degradation rather than by a lack of uptake mechanisms or lack of RNAi machinery activity in midgut tissue.

### Salivary and digestive expression patterns compared to prior H. halys and Nezara viridula studies

Peiffer and Felton (2014) and Liu and Bonning (2019) showed that sheath and watery saliva of *H. halys* harbour distinct protein suites, with watery saliva dominated by principal salivary gland (PSG) proteases and nucleases [11, 12]. This includes a highly-expressed, PSG-derived T2-family nuclease *ribonuclease Oy-like* enzyme in both *H. halys* and the southern green stink bug, *Nezara viridula* [11, 12]. Our atlas confirms these findings by showing that *trypsin* and *chymotrypsin* paralogues, along with a *DNA/RNA non-specific endonuclease* and a *T2-family nuclease*, essentially have salivary gland-specific expression, while cathepsins and aspartic proteases dominate the midgut expression profile. This pattern mirrors the PSG-versus-gut partitioning inferred by Liu and Bonning from three-tissue transcriptomes (PSG, accessory salivary gland, and total gut) and proteomes, and it demonstrates that the same division of enzyme families (serine proteases and nucleases in saliva; cysteine proteases in the gut) persists when considered in a broader, four-organ context that includes the brain/CNS and testes.

Lavore et al. (2018) found that *N. viridula* uses a similar suite of digestive proteases and nucleases, with salivary *ribonuclease Oy-like* and *cathepsins* representing conserved features among phytophagous pentatomids [22]. The near-identical salivary expression of a *ribonuclease Oy-like* gene and the strong midgut *cathepsin L* expansion in *H. halys* suggest that these digestion and dsRNA-degradation modules are phylogenetically stable across Pentatomidae. Against this background, our identification of highly and almost exclusively salivary gland-expressed proteases and nucleases in *H. halys* provides concrete molecular candidates for disrupting feeding and for attenuating the dsRNase barrier that impedes oral RNAi [44, 67, 14].

### Comparative hemipteran analyses of detoxification superfamilies

Volonté et al. (2022) showed that *H. halys* harbours one of the largest complements of detoxification-related genes among surveyed hemipterans, including expansions in CYP3-clan genes, carboxyl/cholinesterases, and sigma-class GSTs [18]. Our tissue-expression atlas extends these genomic observations by demonstrating strong tissue partitioning of these detoxification systems. CYP3 genes and several carboxyl/hormone esterases were predominantly enriched in the midgut (Fig. S5), supporting a major role in digestive xenobiotic metabolism, whereas CYP2-, CYP4-, and mitochondrial CYP-associated expression was more prominent in testes, suggesting involvement in endogenous reproductive metabolism rather than classical detoxification. In contrast, GSTs showed broader expression across tissues, with particularly strong representation in the midgut and brain, consistent with roles in conjugative detoxification and antioxidant defence. These findings refine the hypothesis proposed by Sparks et al. (2020) and Volonté et al. (2022) that detoxification capacity contributes to polyphagy and xenobiotic tolerance in *H. halys* by identifying the midgut as the principal detoxification hub, while also highlighting tissue-specialized metabolic functions in the brain, salivary glands, and testes [6, 18].

When compared to *N. viridula*, which also exhibits expanded CYP and esterase complements but a smaller number of GST genes, *H. halys* emerges as an extreme within Pentatomidae, combining a high gene copy number with pronounced tissue partitioning [6, 18]. This underscores that detoxification-based resistance evolution in *H. halys* may be both genetically and anatomically complex.

### Neuropeptides and GPCR comparisons

Lavore et al. (2018) combined transcriptomics and neuropeptidomics to catalogue neuropeptide precursors and GPCRs in male and female *N. viridula*, showing that most neuropeptides are brain-enriched and that mature peptides are readily detected in brain extracts [22]. Our *H. halys* atlas converges on a similar architecture: most neuropeptide precursors and nearly all biogenic amine receptors and opsins are confined to the brain or strongly brain-enriched. This reinforces the view that central neuroendocrine hubs are conserved across pentatomids and provides a direct *H. halys* reference against which future neuropeptidomic datasets can be interpreted.

Ahn et al. (2020) functionally characterised six PRXamide GPCRs in *H. halys* (pyrokinin and CAPA receptors), demonstrating their ligand specificity in heterologous assays and measuring their expression in body segments by RT-PCR [21]. Our dataset extends this by quantifying PRXamide receptor expression across four discrete tissues and situating them within the broader GPCR landscape that includes all major neuropeptide, biogenic amine, and adhesion GPCRs. In line with Ahn et al., pyrokinin and CAPA receptors are predominantly neural, but we also detect a subset of neuropeptide receptors in the salivary glands and midgut, consistent with local neuromodulation of feeding and secretion. This tissue-level view complements functional receptor characterisation and aligns with conceptual work by Audsley and Down (2015) and Konopińska et al. (2025), who highlight neuropeptides and their GPCRs as promising, highly specific biopesticide targets [19, 20].

### Tau-based tissue specificity and RNAi target selection

Tau analysis identified 4,076 tissue-specific genes (τ ≥ 0.85), with testes contributing the largest fraction, followed by brain, midgut, and salivary glands (Fig. 10). Within these sets, a small number of genes combined very high expression with near-absolute tissue restriction, including the brain-specific components *odorant binding protein, Neuropeptide-like precursor 1, an aminopeptidase N-like gene,* the extremely abundant salivary sheath/matrix proteins (*streptococcal hemagglutinin-like, BadA-like*), and the midgut-enriched ion channels and GPCRs. This use of Tau formalises a strategy that has proven effective in other systems, where highly expressed, tissue-restricted proteins have proven to be effective RNAi targets in experimental work. For example, proteome-guided screening in a mealybug identified *Ferritin* and *OBP* as top candidates whose dsRNA produced the lowest LC50 values [68], and multi-tissue transcriptomes in the fruit fly *Zeugodacus tau* were used to pinpoint highly expressed, testis-specific kinases whose silencing severely disrupted spermatogenesis and male fertility [69].

The classes of high-τ genes prioritised in this study align closely with experimentally tractable targets in other insects. Highly expressed OBPs and related chemosensory proteins have been successfully knocked down to alter host choice and social behaviour in hemipteran pests [70] and ants [71], while neuropeptide/GPCR signalling modules are increasingly recognised as specific, behaviour- and reproduction-modulating insecticide targets [19, 20, 72, 73]. Likewise, salivary sheath proteins functionally analogous to the *H. halys hemagglutinin-like* and *BadA-like* candidates are essential for feeding in planthoppers: dsRNA or antibody-mediated disruption of sheath proteins such as *Nlsp5* impairs sheath formation and drastically reduces phloem ingestion [74]. Finally, midgut-restricted Tau candidates (cathepsins, transporters, channels) fit the same profile as gut proteases and transporters that have repeatedly yielded strong growth and survival phenotypes upon RNAi in hemipteran pests [75], while the large testes-specific Tau set parallels testis-restricted targets now being exploited for sterile-insect strategies in other taxa [69]. Together, these comparisons indicate that our Tau-based, high-expression filter does not simply describe expression patterns but converges with state-of-the-art target-selection frameworks in insect RNAi and provides a rational, tissue-resolved candidate list for *H. halys*. In future studies, we will test the silencing of these highly expressed genes using RNAi to determine functional loss and assess their potential as targets for RNAi pesticides.

### Conceptual advances and outlook

In *H. halys*, prior genome and transcriptome resources, salivary and gut expression studies, and RNAi functional assays have provided solid baseline for RNAi target design and proof-of-function. The present tissue-resolved atlas advances this trajectory by explicitly linking (i) RNAi machinery and nucleases, (ii) detoxification and digestive modules, and (iii) neuropeptides and GPCRs to discrete adult tissues, and by quantifying tissue specificity with the Tau metric to prioritise targets.

Within this framework, salivary gland-expressed trypsins, chymotrypsins, and nucleases; midgut cathepsins and transporters; testes-enriched *AGO-2* isoforms and spermatogenesis genes; and brain-specific neuropeptides, GPCRs, and OBPs emerge as distinct target classes with different delivery and specificity profiles. This is directly aligned with current thinking on RNAi- and neuropeptide-based control strategies, while providing the tissue-level expression context that was missing from previous *H. halys* resources. Future work should extend this atlas to additional tissues, developmental stages, and single-cell resolution, and combine it with systematic RNAi and ligand-screening experiments to turn these expression-based hypotheses into validated targets for species-selective control.

There are several limitations in the current study that should be considered for further studies. The present atlas is restricted to adult males collected in a single season, and transcriptomic profiles in females, nymphal stages, or under active feeding conditions may differ substantially, particularly for salivary secretory proteins and reproductive hormones. Brain-tissue analyses were performed with two biological replicates following outlier exclusion, limiting statistical power for brain-specific differential expression. The 72 h post-injection time point may miss early or transient transcriptional events following dsRNA exposure. Functional validation of top candidate RNAi targets through bioassays, preferably incorporating nanoparticle-formulated oral delivery systems [76–80] remains essential to translate this transcriptomic resource into practical biocontrol application. Nonetheless, our in-depth characterization of gene expression in this wild-caught population of *H. halys* from southern Germany provides both tissue-specific and population-specific transcriptomic resources and a rich set of characterised target genes for functional pest management strategies [4].

## Author contributions

VPSA conceptualised and executed the work. VPSA and SR conducted bioinformatic and statistical analyses, and KAP provided critical review and input on statistical handling and data visualisation. VPSA and SR visualised the results. VPSA wrote the original manuscript. VPSA procured funding. KAP and SR edited and reviewed the manuscript.

## Acknowledgement

VPSA acknowledges the support provided by the state of Baden-Württemberg through bwHPC. VPSA also thanks the programme “Humboldt Reloaded – research right from the start”, organized by the Department of Academic Affairs at the University of Hohenheim, Germany, for funding the RNA-sequencing costs. KAP acknowledges funding support from the Deutsche Forschungsgemeinschaft through grant PA 2044/2-1 within the SPP 2349 “The Genomic Basis of Evolutionary Innovations”.

## Data availability

Raw sequencing data have been deposited in the NCBI Sequence Read Archive (SRA) under BioProject accession PRJNA1451241.

The supporting data used in this study are provided as supplementary information files within this manuscript.

## Declarations

### Ethics approval and consent to participate

The research reported here did not involve experimentation with human participants or animals. Therefore, there was no need for their consent to participate.

### Consent for publication

The research does not contain any individual person’s data in any form; and all authors have consented to publication. There were no human participants so there is no need for participants to consent to publish.

