## supplementary information files for "A tissue-resolved transcriptomic atlas of adult male *Halyomorpha halys* reveals tissue-specific RNAi machinery and a minimal systemic response to non-specific dsRNA": SI_1.pdf

Table S1 Top 20 highest expressed tissue-specific genes across four tissues derived from Tau analysis.

**>dsRNA-GFP sequence**

TGATCGCGCTTCTCGTTGGGGTCTTTGCTCAGGGCGGACTGGGTGCTCAGGTAGTGGTTGTCGGGCAGCAGCA
CGGGGCCGTCGCCGATGGGGGTGTTCTGCTGGTAGTGGTCGGCGAGCTGCACGCTGCCGTCCTCGATGTTGTG
GCGGATCTTGAAGTTCACCTTGATGCCGTTCTTCTGCTTGTCGGCCATGATATAGACGTTGTGGCTGTTGTAGTTG
TACTCCAGCTTGTGCCCCAGGATGTTGCCGTCCTCCTTGAAGTCGATGCCCTTCAGCTCGATGCGGTTCACCAGG
GTGTCGCCCTCGAACTTCACCTCGGCGCGGGTCTTGTAGTTGCCGTCGTCCTTGAAGAAGATGGTGCGCTCCTG
GACGTAGCCTTCGGGCATGGCGGACTTGAAGAAGTCGTGCTGCTTCATGTGGTCGGGGTAGCGGCTGAAGCAC
TGCACGCCGTA

**Brain replicate exclusion criteria**

During gene-family–focused expression profiling, one brain biological replicate from each treatment group (replicate 1 in both UNT and dsGFP conditions) exhibited anomalous expression patterns relative to the other replicates within the same group. These deviations were most evident within specific gene families, including nucleases and proteases, where replicate 1 consistently showed discordant normalized expression profiles.

Importantly, this discrepancy was not detected in principal component analysis (PCA) of global variance-stabilized counts, indicating that the effect was not driven by a global transcriptomic shift but was instead restricted to specific subsets of genes.

Consistent with this observation, the heatmap of the top variable genes (Fig. S1) reveals that brain replicate 1 does not cluster with the other brain replicates. Instead, the remaining brain replicates within each treatment group cluster together and display concordant expression patterns, whereas replicate 1 shows clear divergence across multiple gene clusters.

Based on these reproducible, gene-family–specific inconsistencies and its distinct clustering behavior, these samples were classified as outliers and excluded from downstream analyses. Consequently, brain tissue analyses were conducted using n = 2 biological replicates per treatment group, while all other tissues (midgut, salivary glands, and testes) retained n = 3 biological replicates per treatment group.

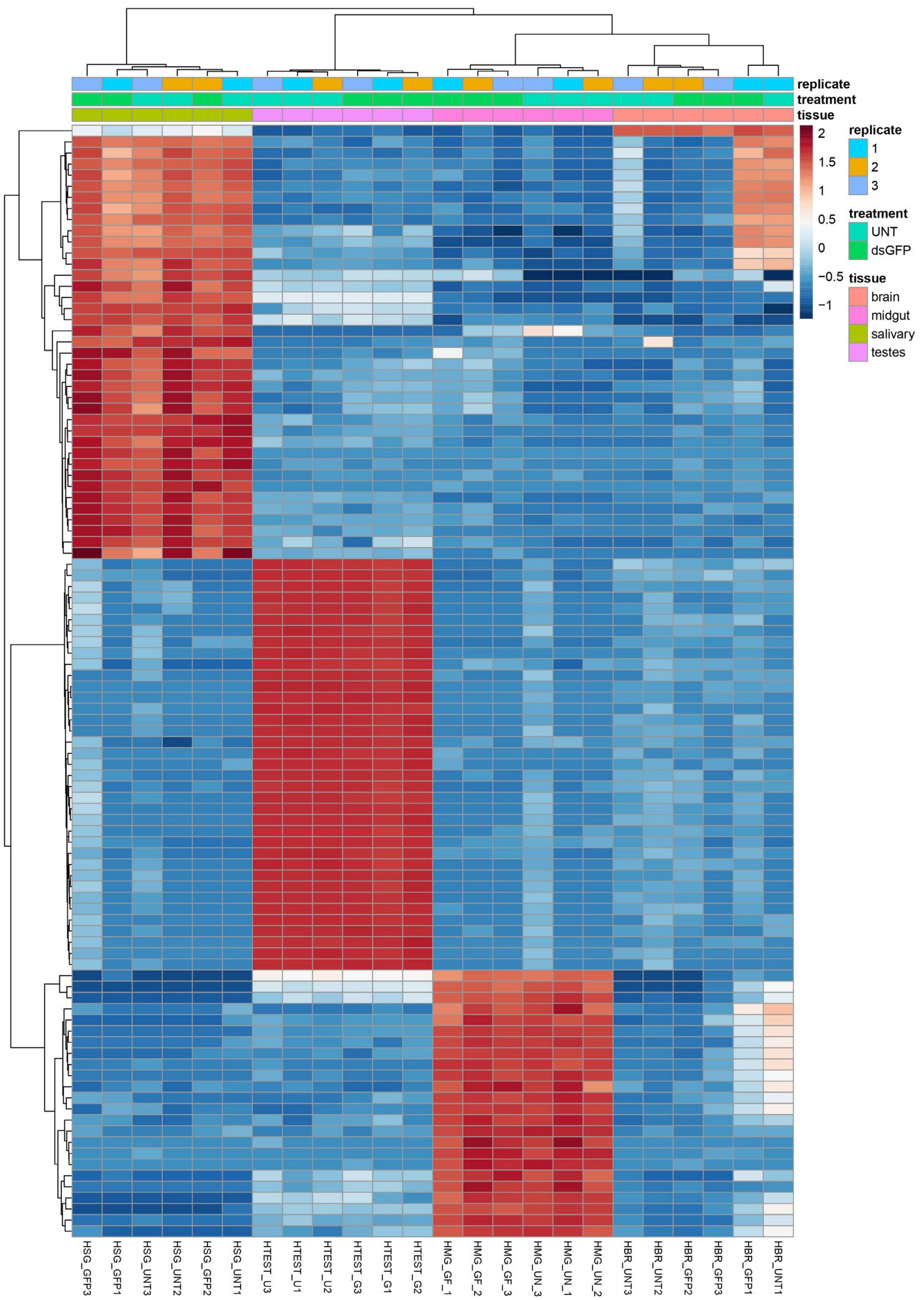

*Figure S1* ***Replicate-level heatmap of the most variable genes across tissues and treatments in Halyomorpha halys.***

*Heatmap showing variance-stabilized expression values (DESeq2 VST) for the top 100 most variable genes across all biological replicates. Rows represent genes and columns represent individual samples. Expression values are row-scaled (z-scores), with red indicating higher relative expression and blue indicating lower relative expression. Column annotations indicate biological replicate, treatment group (dsGFP = dsRNA-GFP injected group or UNT = nuclease-free water injected group), and tissue type (brain, midgut, salivary glands, and testes). Hierarchical clustering was performed using Euclidean distance and complete linkage. Brain replicates 1 from both treatment groups displays distinct clustering behaviour relative to the other brain replicates, which group together and exhibit more consistent expression patterns. This divergence is consistent across multiple gene clusters and supports its classification as an outlier. In contrast, replicates from other tissues show tighter clustering within their respective groups. This heatmap supports the identification and exclusion of anomalous brain replicates based on gene-level expression inconsistencies rather than global transcriptomic variation.*

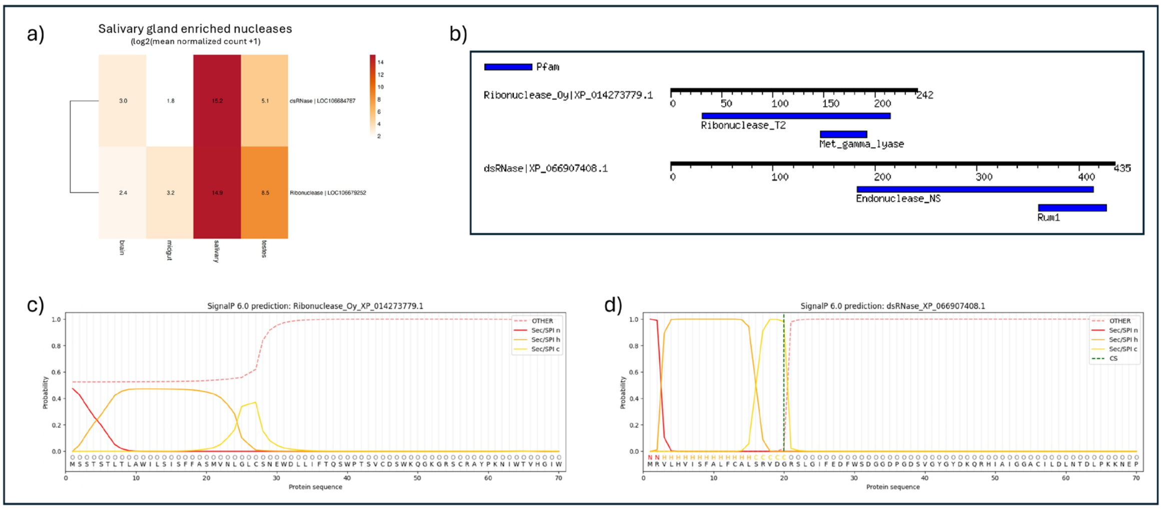

*Figure S2* ***Characterisation of salivary gland-enriched nuclease candidates.***

*(A) Heatmap showing $\log_2$(mean normalised count + 1) expression values of two nuclease genes across brain, midgut, salivary gland, and testes tissues. Rows represent genes and columns represent tissues; colour intensity indicates relative expression level from low (light) to high (dark). (B) Protein domain architectures of Ribonuclease_Oy (XP_014273779.1) and dsRNase (XP_066907408.1). Black horizontal lines indicate protein length in amino acids, and blue boxes indicate predicted conserved domains annotated by Pfam. Amino acid positions are shown above each protein schematic. (C–D) SignalP 6.0 predictions for Ribonuclease_Oy (C) and dsRNase (D). The x-axis represents amino acid position within the protein sequence and the y-axis indicates prediction probability. Coloured lines correspond to SignalP prediction classes: SP (signal peptide), CS (cleavage site), OTHER, Sec/SPI, and Sec/SPII*

Figure S3. **Tissue-specific expression profiles of annotated nuclease-related genes.**

*Heatmap showing log2(mean normalised read count + 1) expression values of predicted nuclease-related genes across four tissues: brain, midgut, salivary gland, and testes. Rows represent individual genes and columns represent tissues. Gene names and locus identifiers are listed at right. Numbers within cells indicate expression values. Colour intensity ranges from low expression (light yellow) to high expression (dark red), as shown by the scale bar. Coloured side bars indicate functional annotation categories: DNase, phosphatase/nucleotidase, RNA metabolism regulator, RNA-processing complex, RNase, and RNase-related. Genes are grouped by functional class and separated by white horizontal lines.*

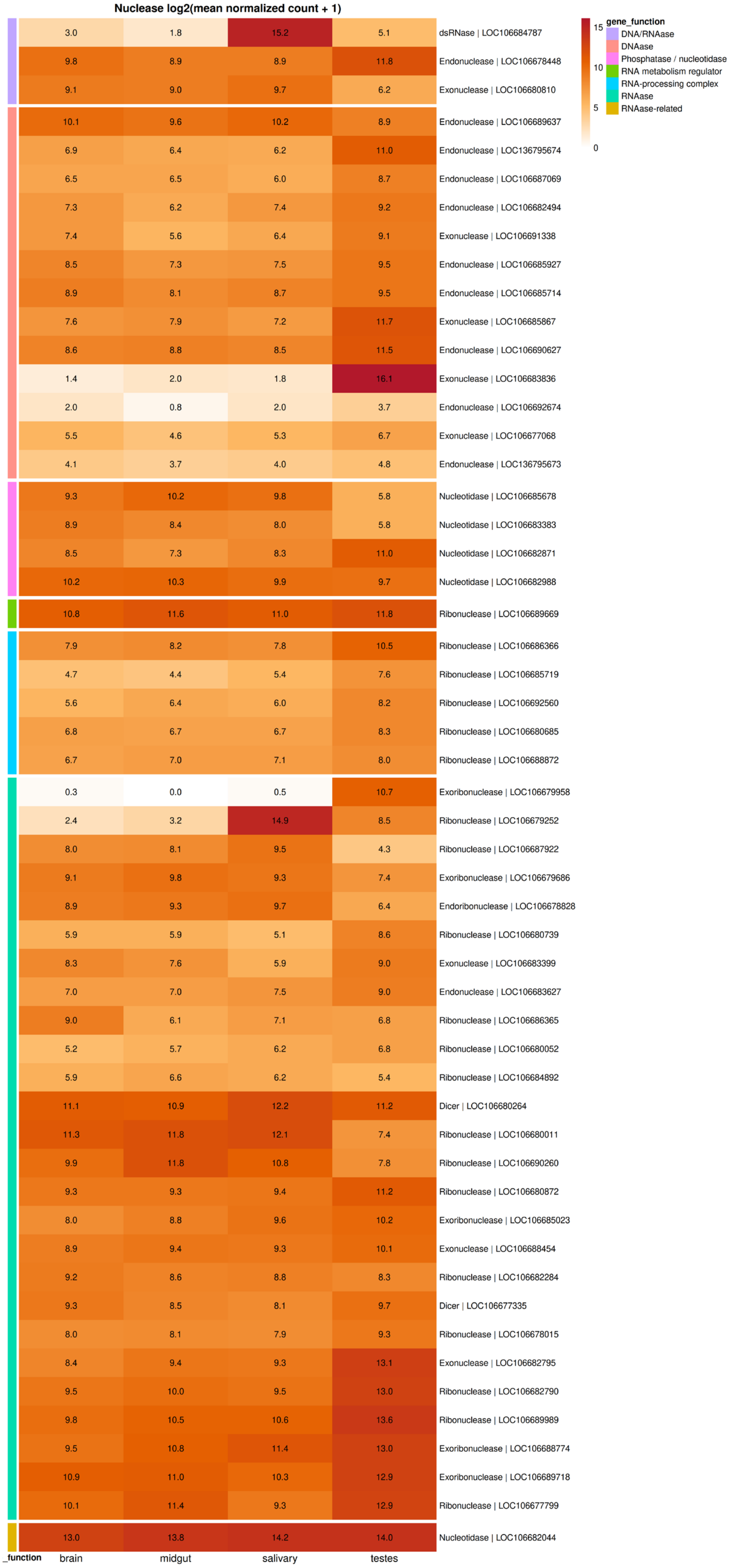

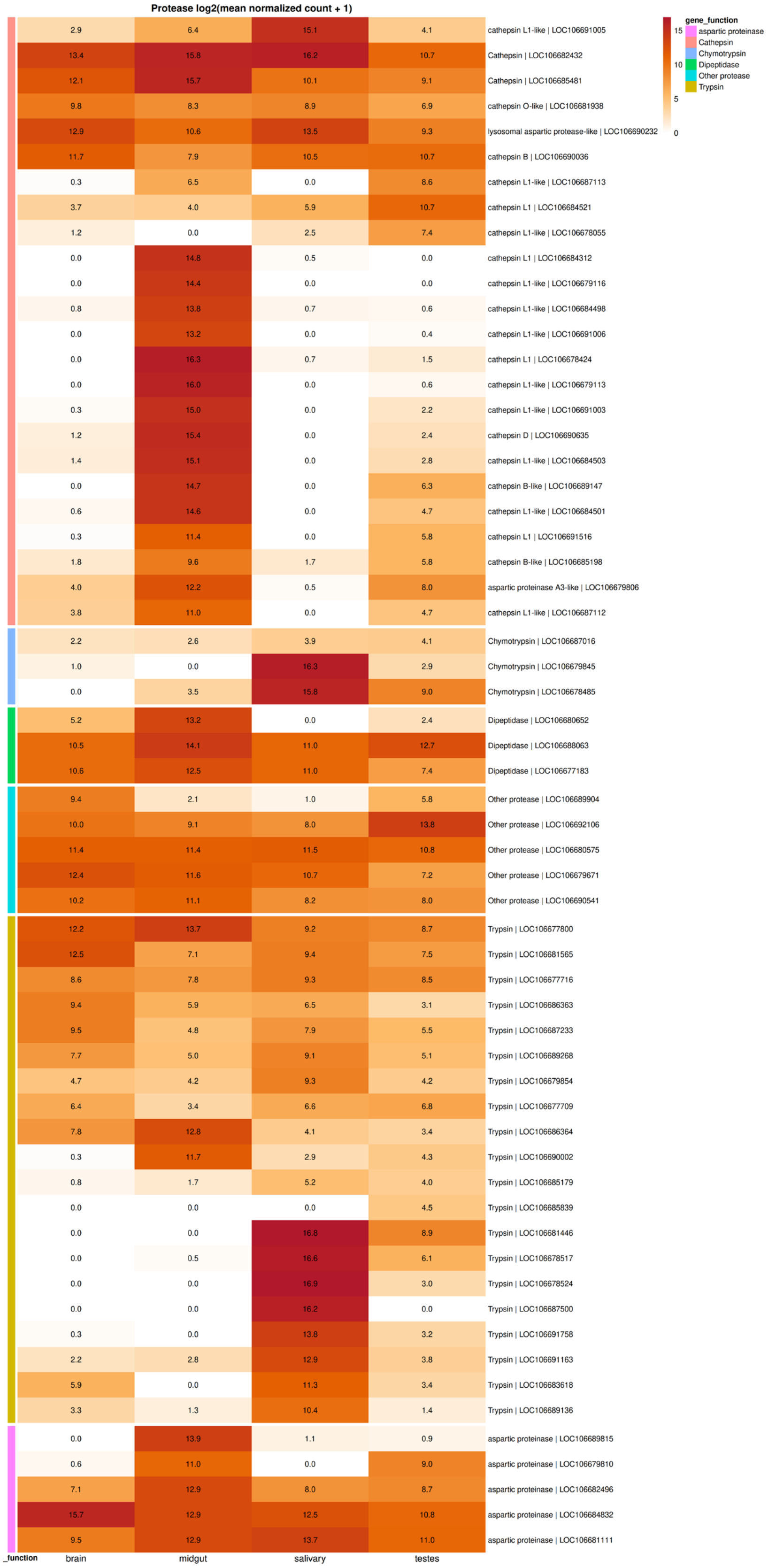

Figure S4 **Tissue-specific expression profiles of annotated protease genes.**

*Heatmap showing log2(mean normalised read count + 1) expression values of predicted protease genes across four tissues: brain, midgut, salivary gland, and testes. Rows represent individual genes and columns represent tissues. Gene names and locus identifiers are listed at right. Numbers within cells indicate expression values. Colour intensity ranges from low expression (light grey/yellow) to high expression (dark red), as shown by the scale bar. Coloured side bars indicate functional annotation categories: aspartic protease, cathepsin, chymotrypsin, dipeptidase, other protease, and trypsin. Genes are grouped by functional class and separated by white horizontal lines.*

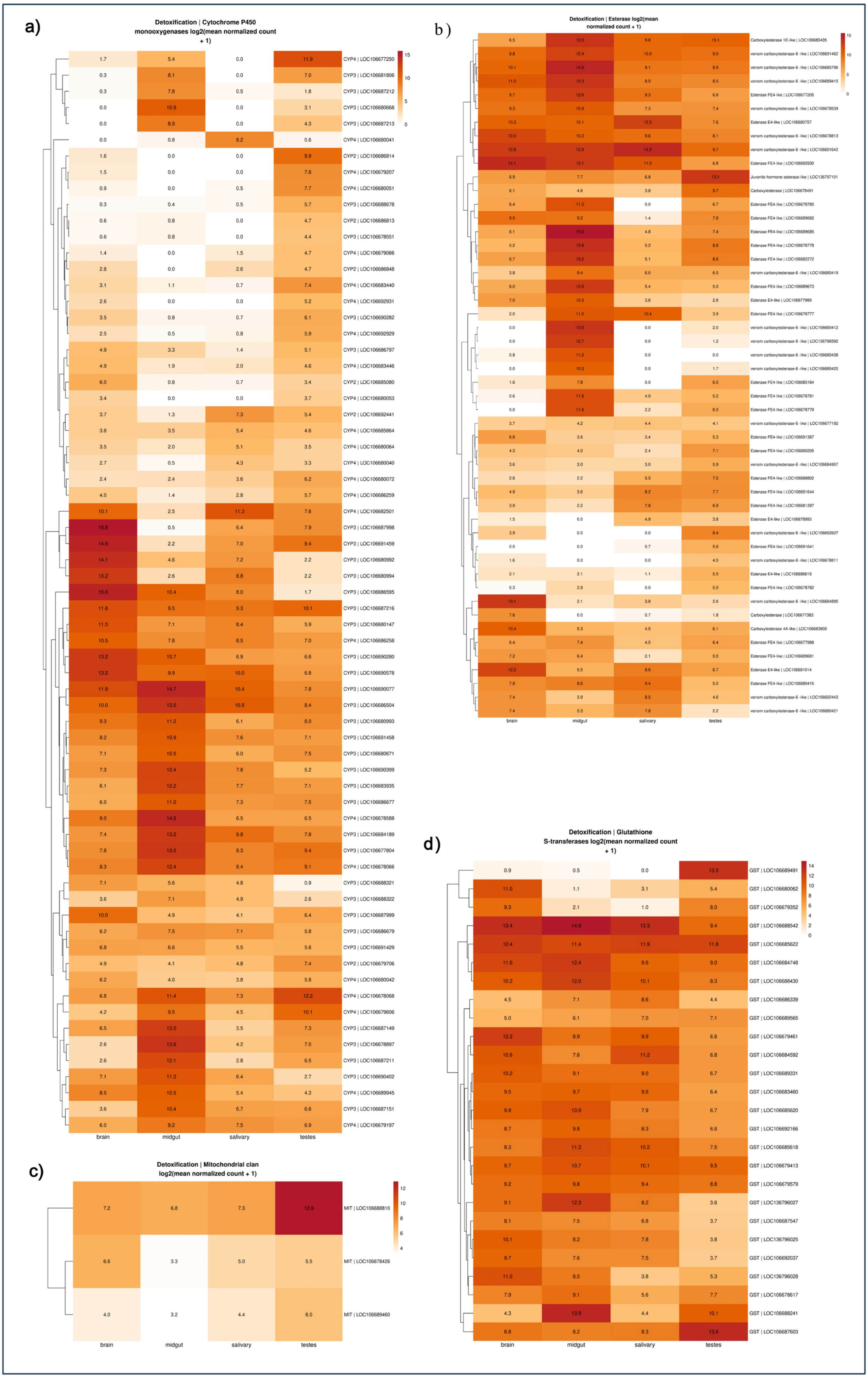
a)
Detoxification | Cytochrome P450 monooxygenases log2(mean normalized count + 1)
b)
Detoxification | Esterase log2(mean normalized count + 1)
c)
Detoxification | Mitochondrial clan log2(mean normalized count + 1)
d)
Detoxification | Glutathione S-transferases log2(mean normalized count + 1)
brain
midgut
salivary
testes

*Figure S5* ***Tissue-resolved expression patterns of detoxification-associated gene families.***

*Heatmaps showing normalised expression values of detoxification-related genes across tissues. (A) Cytochrome P450 monooxygenases (CYPs). (B) Esterases and carboxylesterases. (C) Mitochondrial clan CYP genes. (D) Glutathione S-transferases (GSTs). Rows represent individual genes clustered by expression similarity, and columns represent tissues. The colour scale indicates expression intensity from low (light) to high (dark).*

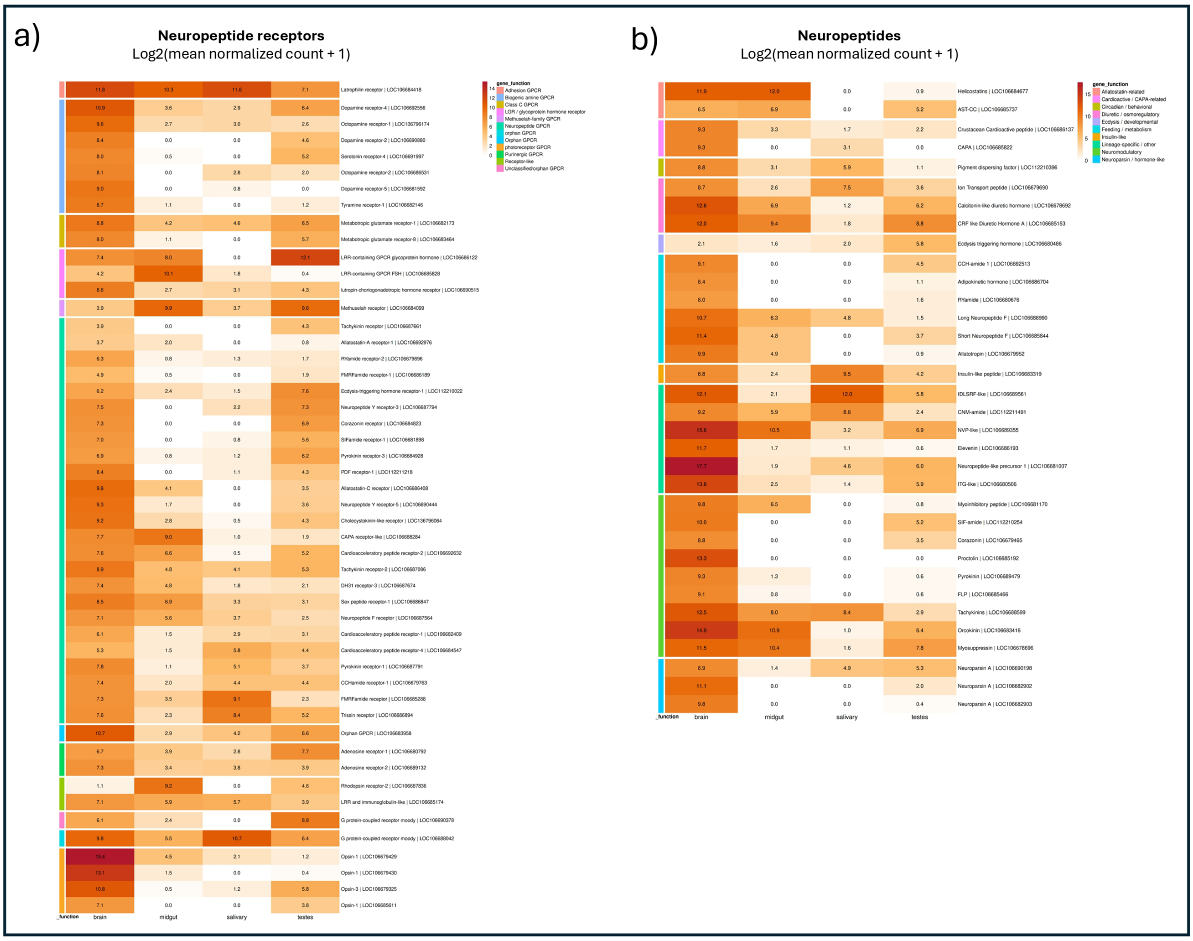

*Figure S6* ***Tissue-resolved expression patterns of neuropeptide receptors and neuropeptides.***

*(a) Neuropeptide receptors. (b) Neuropeptides. Heatmaps show normalised expression values. Rows represent individual genes grouped by functional categories and clustered by expression similarity, and columns represent tissues. The colour scale indicates expression intensity from low (light) to high (dark).*

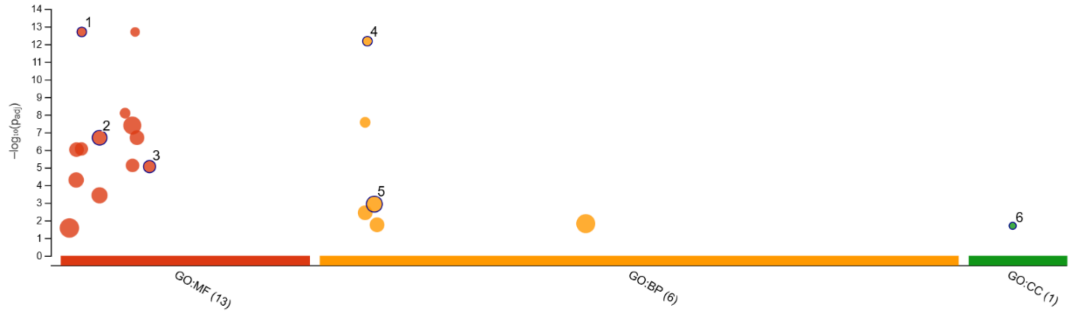

| ID | Source | Term ID | Term Name | $p_{adj}$ (query_1) |
|---|---|---|---|---|
| 1 | GO:MF | GO:0004567 | beta-mannosidase activity | $2.009\times10^{-13}$ |
| 2 | GO:MF | GO:0008236 | serine-type peptidase activity | $2.019\times10^{-7}$ |
| 3 | GO:MF | GO:0030246 | carbohydrate binding | $8.739\times10^{-6}$ |
| 4 | GO:BP | GO:0006080 | substituted mannan metabolic process | $6.733\times10^{-13}$ |
| 5 | GO:BP | GO:0006508 | proteolysis | $1.182\times10^{-3}$ |
| 6 | GO:CC | GO:0042721 | TIM22 mitochondrial import inner membrane inse... | $2.006\times10^{-2}$ |

**version** e113_eg59_p19_6be52918
**date** 3/22/2026, 5:33:47 PM
**organism** hhgca000696795v2rs

g:Profiler

*Figure S7 Salivary gland enriched genes – gProfiler GO enrichment analysis*

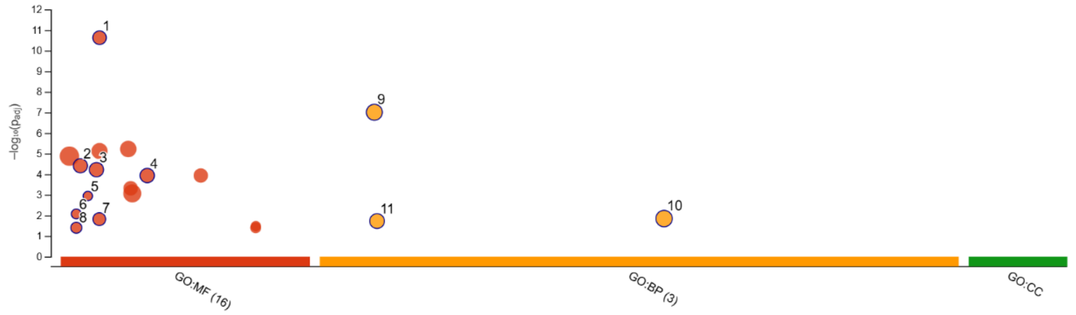

| ID | Source | Term ID | Term Name | $p_{adj}$ (query_1) |
|---|---|---|---|---|
| 1 | GO:MF | GO:0008234 | cysteine-type peptidase activity | $2.344\times10^{-11}$ |
| 2 | GO:MF | GO:0004497 | monooxygenase activity | $3.871\times10^{-5}$ |
| 3 | GO:MF | GO:0005506 | iron ion binding | $6.012\times10^{-5}$ |
| 4 | GO:MF | GO:0020037 | heme binding | $1.156\times10^{-4}$ |
| 5 | GO:MF | GO:0004869 | cysteine-type endopeptidase inhibitor activity | $1.142\times10^{-3}$ |
| 6 | GO:MF | GO:0004185 | serine-type carboxypeptidase activity | $8.474\times10^{-3}$ |
| 7 | GO:MF | GO:0008194 | UDP-glycosyltransferase activity | $1.513\times10^{-2}$ |
| 8 | GO:MF | GO:0004190 | aspartic-type endopeptidase activity | $3.960\times10^{-2}$ |
| 9 | GO:BP | GO:0006508 | proteolysis | $9.812\times10^{-8}$ |
| 10 | GO:BP | GO:0055085 | transmembrane transport | $1.419\times10^{-2}$ |
| 11 | GO:BP | GO:0006629 | lipid metabolic process | $1.883\times10^{-2}$ |

**version** e113_eg59_p19_6be52918
**date** 3/22/2026, 5:32:58 PM
**organism** hhgca000696795v2rs

g:Profiler

*Figure S8 midgut enriched genes – gProfiler GO enrichment analysis*

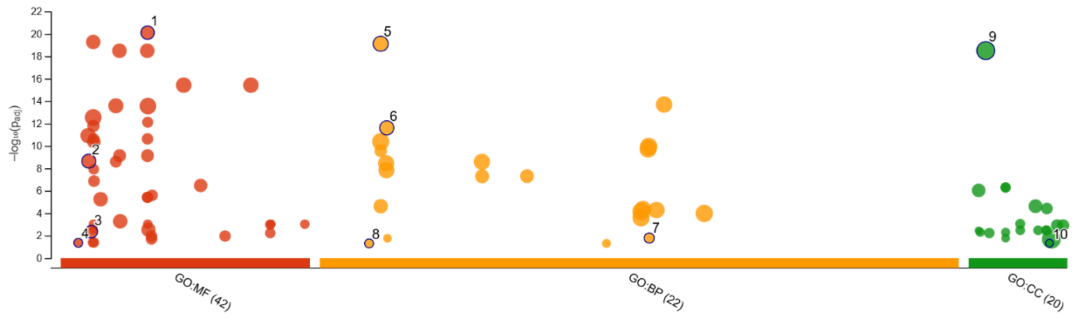

| ID | Source | Term ID | Term Name | $p_{adj}$ (query_1) |
|---|---|---|---|---|
| 1 | GO:MF | GO:0022836 | gated channel activity | $7.828\times10^{-21}$ |
| 2 | GO:MF | GO:0004930 | G protein-coupled receptor activity | $2.269\times10^{-9}$ |
| 3 | GO:MF | GO:0005102 | signaling receptor binding | $4.393\times10^{-3}$ |
| 4 | GO:MF | GO:0004383 | guanylate cyclase activity | $4.353\times10^{-2}$ |
| 5 | GO:BP | GO:0006811 | monoatomic ion transport | $7.637\times10^{-20}$ |
| 6 | GO:BP | GO:0007186 | G protein-coupled receptor signaling pathway | $2.419\times10^{-12}$ |
| 7 | GO:BP | GO:0051260 | protein homooligomerization | $1.643\times10^{-2}$ |
| 8 | GO:BP | GO:0006182 | cGMP biosynthetic process | $4.978\times10^{-2}$ |
| 9 | GO:CC | GO:0016020 | membrane | $3.111\times10^{-19}$ |
| 10 | GO:CC | GO:0099503 | secretory vesicle | $4.682\times10^{-2}$ |

**version** e113_eg59_p19_6be52918
**date** 3/22/2026, 5:28:57 PM
**organism** hhgca000696795v2rs

g:Profiler

*Figure S9 Brain/CNS enriched genes – gProfiler GO enrichment analysis*

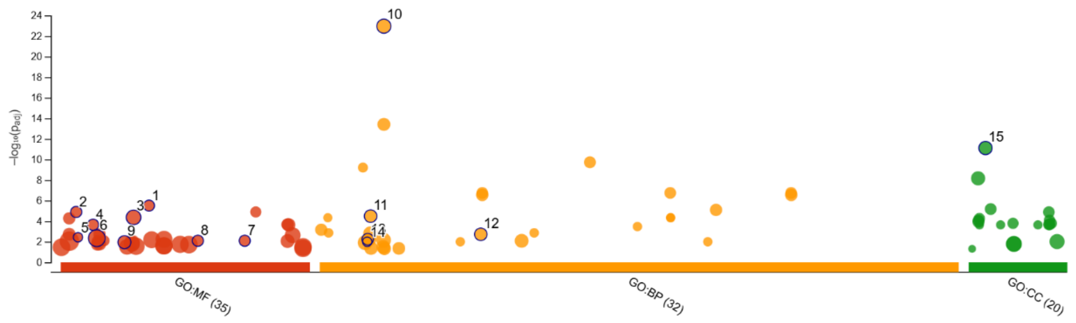

| ID | Source | Term ID | Term Name | $p_{adj}$ (query_1) |
|---|---|---|---|---|
| 1 | GO:MF | GO:0030145 | manganese ion binding | $3.122\times10^{-6}$ |
| 2 | GO:MF | GO:0004177 | aminopeptidase activity | $1.294\times10^{-5}$ |
| 3 | GO:MF | GO:0016853 | isomerase activity | $4.357\times10^{-5}$ |
| 4 | GO:MF | GO:0005200 | structural constituent of cytoskeleton | $2.271\times10^{-4}$ |
| 5 | GO:MF | GO:0004371 | glycerone kinase activity | $3.719\times10^{-3}$ |
| 6 | GO:MF | GO:0005524 | ATP binding | $4.579\times10^{-3}$ |
| 7 | GO:MF | GO:0051959 | dynein light intermediate chain binding | $8.126\times10^{-3}$ |
| 8 | GO:MF | GO:0045505 | dynein intermediate chain binding | $8.126\times10^{-3}$ |
| 9 | GO:MF | GO:0015631 | tubulin binding | $1.128\times10^{-2}$ |
| 10 | GO:BP | GO:0007017 | microtubule-based process | $1.099\times10^{-23}$ |
| 11 | GO:BP | GO:0006260 | DNA replication | $3.288\times10^{-5}$ |
| 12 | GO:BP | GO:0022402 | cell cycle process | $1.924\times10^{-3}$ |
| 13 | GO:BP | GO:0006099 | tricarboxylic acid cycle | $5.735\times10^{-3}$ |
| 14 | GO:BP | GO:0006071 | glycerol metabolic process | $1.022\times10^{-2}$ |
| 15 | GO:CC | GO:0015630 | microtubule cytoskeleton | $7.810\times10^{-12}$ |

**version** e113_eg59_p19_6be52918
**date** 3/22/2026, 5:34:29 PM
**organism** hhgca000696795v2rs

g:Profiler

*Figure S10 Testes enriched genes – gProfiler GO enrichment analysis*

*Table S1* ***Top 20 highest expressed tissue-specific genes across four tissues derived from Tau analysis.***

*Top 20 highest expressed genes classified as tissue-specific in brain, midgut, salivary glands, and testes based on control-only CPM-normalised expression. Tissue specificity was defined using the Tau (τ) metric (τ ≥ 0.85) with expression thresholds (CPM ≥ 10 and ≥ 5% of maximum expression) applied. Values represent mean expression per tissue. Genes are ordered by decreasing expression within each tissue and represent candidate tissue-restricted functional genes.*

| gene_id | gene_name | brain | midgut | salivary | testes | tau | tissue |
|---|---|---|---|---|---|---|---|
| LOC106688070 | odorant binding protein | 10375.15388 | 0.022760108 | 15.20840914 | 5.24820097 | 0.999342038 | brain |
| LOC106681007 | neuropeptide-like 1 | 6711.277135 | 0.084289502 | 0.708604861 | 1.693300436 | 0.999876517 | brain |
| LOC106686134 | aminopeptidase N-like | 4182.36322 | 0.16991547 | 3.774910966 | 2.791071272 | 0.999463151 | brain |
| LOC106690807 | uncharacterized | 3625.593345 | 0.011380054 | 12.04270102 | 0.006844596 | 0.998891131 | brain |
| LOC106678115 | sodium/potassium-transporting ATPase subunit alpha | 3383.108738 | 131.9812857 | 102.8219056 | 11.49361005 | 0.975732694 | brain |
| LOC106679216 | uncharacterized | 2895.718403 | 0.011380054 | 1.303634435 | 0.063605027 | 0.999841304 | brain |
| LOC106684983 | glutamate-gated chloride channel | 2509.91188 | 0.252818999 | 0.289811523 | 1.683306719 | 0.99970438 | brain |
| LOC106685454 | uncharacterized | 2215.253039 | 23.81195231 | 0 | 0.795479453 | 0.996297273 | brain |
| LOC106680960 | inhibin beta E chain | 1983.612732 | 0 | 0.024681758 | 0.050395746 | 0.999987384 | brain |
| LOC106687998 | cytochrome P450 3A25-like | 1816.832996 | 0.011380054 | 2.685658277 | 5.952416252 | 0.998413089 | brain |
| LOC106681727 | serine/threonine-protein phosphatase 6 regulatory ankyrin repeat subunit C-like | 1582.464135 | 0.216221029 | 14.43358164 | 0.055977547 | 0.996902346 | brain |
| LOC106689355 | uncharacterized | 1517.868122 | 43.87423558 | 0.245535696 | 3.048884068 | 0.989641468 | brain |
| LOC106681741 | serine/threonine-protein phosphatase 6 regulatory ankyrin repeat subunit A-like | 1405.416924 | 0.534584094 | 28.14937132 | 0.071889354 | 0.993179759 | brain |
| LOC106677204 | mucin-5AC-like | 1398.326755 | 0.036454724 | 0.638607073 | 2.162854663 | 0.999323497 | brain |
| LOC106679429 | opsin-1 | 1388.11494 | 0.63412784 | 0.123408792 | 0.035224811 | 0.999809631 | brain |
| LOC106683425 | receptor-type tyrosine-protein phosphatase N2 | 1334.0199 | 4.885534549 | 0.061704396 | 13.68445162 | 0.995344475 | brain |
| LOC106684106 | sodium/potassium-transporting ATPase subunit beta-2-like | 1294.421214 | 0.438350975 | 4.915413524 | 0.843451777 | 0.998404122 | brain |
| LOC136795769 | Tyramine beta hydroxylase | 1287.700115 | 0.105963953 | 0.132700561 | 0.889594774 | 0.999707939 | brain |
| LOC106687738 | ankyrin-3-like | 1232.553311 | 0 | 0 | 0.183709525 | 0.999950317 | brain |
| LOC106690802 | lncRNA | 1208.646711 | 0.703949823 | 0.024681758 | 6.355297762 | 0.998046319 | brain |
| LOC106682723 | soma ferritin | 12.05869185 | 61551.76888 | 268.8552844 | 2.326387967 | 0.998466113 | midgut |
| LOC106692576 | uncharacterized | 0 | 11463.39136 | 0 | 9.439524523 | 0.999725517 | midgut |
| LOC106687253 | uricase | 0.025408484 | 11192.12074 | 0.056859373 | 0.020533787 | 0.999996938 | midgut |
| LOC106683950 | venom carboxylesterase-6-like | 0.130970645 | 10005.97644 | 0 | 2.702906983 | 0.999905594 | midgut |
| LOC106679058 | WSC domain-containing protein ARB_07867 | 0.970873594 | 7399.579041 | 4.631850024 | 1.93044508 | 0.999660649 | midgut |
| LOC106683269 | uncharacterized | 0 | 7395.179047 | 0 | 0 | 1 | midgut |
| LOC112210696 | uncharacterized | 0.162956428 | 7185.590988 | 0 | 0.228607496 | 0.999981836 | midgut |
| LOC106685944 | uncharacterized | 0.098227984 | 7057.748416 | 0.018953124 | 1.532012141 | 0.99992211 | midgut |
| LOC106686132 | aminopeptidase N-like | 0.025030045 | 7030.152342 | 0 | 0.084921699 | 0.999994787 | midgut |
| LOC106684321 | cathepsin L1 | 0 | 6995.169421 | 0.012340879 | 0 | 0.999999412 | midgut |
| LOC106685942 | uncharacterized | 0.049303211 | 6646.604299 | 0.13267187 | 0.647172048 | 0.999958418 | midgut |
| LOC106687106 | xylosyltransferase oxt | 0.024273167 | 6572.255294 | 0 | 0.988052786 | 0.999948657 | midgut |
| LOC106681980 | basic proline-rich protein-like | 0.008469495 | 6200.759122 | 0.056859373 | 0.177908775 | 0.999986924 | midgut |
| LOC106690764 | lipase 3-like | 2.8937896 | 6025.16026 | 0.037906248 | 0.589409783 | 0.9998052 | midgut |
| LOC106681981 | basic proline-rich protein-like | 0 | 5862.898867 | 0 | 0 | 1 | midgut |
| LOC106685659 | uncharacterized | 0.008469495 | 5555.545742 | 0.018953124 | 0.111862184 | 0.999991643 | midgut |

| | | | | | | | |
|---|---|---|---|---|---|---|---|
| LOC106680650 | uncharacterized | 0.033499539 | 5375.873962 | 0.0929984 | 0.006844596 | 0.999991732 | midgut |
| LOC106678109 | uncharacterized | 0.024273167 | 5116.348023 | 0.012340879 | 0.338483968 | 0.999975562 | midgut |
| LOC106679110 | cathepsin L1-like | 0.008091056 | 5054.857696 | 0 | 0.189636313 | 0.999986961 | midgut |
| LOC106692511 | uncharacterized | 0.073576378 | 4990.703062 | 0.111951524 | 0.605456513 | 0.999947169 | midgut |
| LOC106690997 | streptococcal hemagglutinin | 0.008091056 | 0.131274226 | 224615.1948 | 0.062822143 | 0.9999997 | salivary |
| LOC106690998 | streptococcal hemagglutinin-like | 0.01656055 | 0.035604669 | 159146.3785 | 0.217587516 | 0.999999435 | salivary |
| LOC106690607 | autotransporter adhesin BadA | 0.514746971 | 0.525755586 | 37027.63486 | 7.778307369 | 0.999920611 | salivary |
| LOC106679835 | uncharacterized | 0 | 0 | 36741.45117 | 0.014169101 | 0.999999871 | salivary |
| LOC106690464 | keratinocyte proline-rich protein | 0.056637389 | 0.02260304 | 34923.71062 | 3.791928571 | 0.999963051 | salivary |
| LOC106690702 | oleosin-B6 | 0.041212156 | 0.161164114 | 34325.85343 | 0.928691623 | 0.999989016 | salivary |
| LOC106686694 | streptococcal hemagglutinin | 0.0331211 | 0 | 31756.82452 | 0.254033109 | 0.999996986 | salivary |
| LOC106682756 | uncharacterized | 0 | 0 | 28668.19128 | 5.440155776 | 0.999936746 | salivary |
| LOC136796531 | mucin-2-like | 0 | 0 | 27661.42357 | 0.064825809 | 0.999999219 | salivary |
| LOC106685636 | vasotab-like | 1.477870259 | 0.106042487 | 27540.67792 | 2.93669651 | 0.999945286 | salivary |
| LOC106691110 | uncharacterized lncRNA | 0 | 0.047063258 | 18012.71087 | 0.085317583 | 0.99999755 | salivary |
| LOC106690127 | spidroin-1-like | 3.836674096 | 0.113643473 | 15946.85114 | 2.164720439 | 0.999872179 | salivary |
| LOC106682271 | uncharacterized | 8.74661952 | 9.613251295 | 13054.71621 | 0.828574932 | 0.999510051 | salivary |
| LOC106691348 | spidroin-1 | 3.974222041 | 0.071287873 | 11034.43458 | 0.163268648 | 0.999872859 | salivary |
| LOC106688561 | uncharacterized protein | 0 | 0.011380054 | 9927.633543 | 0.621437171 | 0.999978752 | salivary |
| LOC106686960 | putative mediator of RNA polymerase II transcription subunit 15 | 0.140953895 | 0 | 9915.493435 | 0.126031286 | 0.999991025 | salivary |
| LOC106685394 | xanthine dehydrogenase/oxidase | 31.34385394 | 0.324342475 | 8678.739884 | 38.83240781 | 0.99729221 | salivary |
| LOC106687558 | uncharacterized | 0 | 0 | 8177.745009 | 0.098568877 | 0.999995982 | salivary |
| LOC106679471 | K+-dependent Na+/Ca+ exchanger | 3.495336006 | 0.056900271 | 7956.908562 | 0.013689191 | 0.999850615 | salivary |
| LOC106679217 | uro-adherence factor A | 0 | 0.207377317 | 7536.733285 | 5.905154112 | 0.999729656 | salivary |
| LOC106688998 | methenyltetrahydrofolate synthase domain-containing protein lost | 0.624615798 | 0.714464618 | 0.055975762 | 7614.430299 | 0.999938929 | testes |
| LOC106679860 | serine-rich adhesin for platelets | 0.09316444 | 0.113800542 | 0.049363517 | 4846.232473 | 0.999982369 | testes |
| LOC106679128 | uncharacterized | 0.140953895 | 0.034140163 | 0 | 4654.835334 | 0.999987461 | testes |
| LOC106682092 | outer dense fiber protein 2 | 1.096780695 | 0.207077002 | 0.156498709 | 4478.281568 | 0.999891301 | testes |
| LOC106682308 | tubulin beta-4B chain | 0.100498617 | 0.079660379 | 0 | 4396.205098 | 0.99998634 | testes |
| LOC106679052 | Tubulin polyglutamylase TTLL5 | 0.376585986 | 0.147233896 | 0.111067913 | 3885.586758 | 0.999945535 | testes |
| LOC106684175 | cytosol aminopeptidase-like | 0.09316444 | 0.034140163 | 0.082453434 | 3679.14709 | 0.999980996 | testes |
| LOC106684618 | cytosol aminopeptidase-like | 0.250300446 | 0.079503311 | 0.012340879 | 3643.066419 | 0.999968694 | testes |
| LOC106686754 | hornerin-like | 0.20908829 | 0.045520217 | 0.024681758 | 3569.910386 | 0.999973922 | testes |
| LOC106678194 | uncharacterized | 0.090893806 | 0.034140163 | 0.17365592 | 3542.098156 | 0.999971891 | testes |
| LOC106690857 | putative leucine-rich repeat-containing protein | 2.184335018 | 0.116115103 | 0.275946089 | 3435.678986 | 0.999750035 | testes |
| LOC106691735 | protein FAM133 | 0.109346551 | 0.102420488 | 0.012340879 | 3295.475763 | 0.999977332 | testes |
| LOC106686346 | uncharacterized | 0.09316444 | 0.034140163 | 0.012340879 | 3177.247918 | 0.999985349 | testes |
| LOC106680018 | cytosol aminopeptidase-like | 0.124393345 | 0.045520217 | 0 | 3030.680832 | 0.999981312 | testes |
| LOC106688226 | heat shock protein 68 | 0.332346319 | 0.183074168 | 0.065908475 | 2989.646042 | 0.999935184 | testes |
| LOC106692802 | tubulin alpha-4 chain | 0.125528662 | 0.13771102 | 0.012340879 | 2897.128068 | 0.999968293 | testes |
| LOC106677619 | glutamate dehydrogenase, mitochondrial | 0.073954817 | 0.125180596 | 0.094794313 | 2670.527541 | 0.999963312 | testes |
| LOC106689448 | uncharacterized | 0.484562619 | 0.045520217 | 0.012340879 | 2631.214398 | 0.999931283 | testes |

| | | | | | | | |
|---|---|---|---|---|---|---|---|
| LOC106681927 | MATH and LRR domain-containing protein PFE0570w-like | 0.500366291 | 0.09173342 | 0.908651398 | 2621.103591 | 0.999809145 | testes |
| LOC106677633 | uncharacterized | 0.073576378 | 0.022760108 | 0.024681758 | 2578.939578 | 0.999984358 | testes |
